# MintCNA: A Unified Framework for Integrative Copy Number Profiling with Single-Cell Multi-Omics Data

**DOI:** 10.64898/2026.06.26.734559

**Authors:** Wenhan Bao, Feifei Xiao

**Affiliations:** Department of Biostatistics, College of Public Health and Health Professions and College of Medicine, University of Florida, Gainesville, FL, 32603, USA

## Abstract

Chromosomal copy number alterations (CNAs) are key drivers of tumor evolution, disease progression and therapeutic resistance, and their identification is critical for delineating tumor clonal structure. However, accurately detecting CNAs from single-cell data remains challenging as most existing CNA callers still analyze a single omics layer and are susceptible to assay-specific noise. Recently, single-cell multi-omics techniques have enabled joint sequencing of multiple molecular layers (e.g., genomics and transcriptomics) simultaneously in the same cells, yet computational methods that fully exploit such complementary multi-omics data for CNA analysis remain limited. Here, we present MintCNA, a unified framework for CNA detection from paired single-cell multi-omics data. MintCNA integrates deep-learning based denoising with a missingness-adjusted multivariate change-point detection procedure to enhance CNA profiling across omics layers. In simulations, MintCNA outperformed existing single-cell CNA callers across all evaluated scenarios. MintCNA showed the highest correlation with bulk whole-genome sequencing in a colorectal cancer dataset and revealed progressive loss of chromosome 17 during metastasis. In an IDH-mutant glioma cohort, MintCNA generalized to paired single-cell ATAC-seq and RNA-seq data and identified copy number subclones corresponding to distinct malignant cell states. MintCNA provides an advanced multi-omics integrative single-cell CNA analysis tool for better delineating the mechanisms underlying disease progression and tumor evolution.

## Introduction

Cancer genomes carry various types of somatic mutations, including single nucleotide variants and copy number alterations (CNAs) (Stratton 2011; Vogelstein et al. 2013). CNAs are deletions or duplications of genomic segments that accumulate progressively during tumor evolution, promoting tumor growth by disrupting the functions of oncogenes and tumor suppressors (Zack et al. 2013). For example, amplification of the oncogene *MYC* at 8q24 is recurrent in breast and colorectal cancers, while homozygous deletion of the tumor suppressor *CDKN2A* at 9p21 is prevalent in melanoma and pancreatic cancer (Beroukhim et al. 2010). Driven in part by CNAs, cancer cells diverge into distinct clonal subpopulations, and this intratumor heterogeneity has been associated with metastatic progression and treatment resistance (Jamal-Hanjani et al. 2015). Therefore, accurate CNA profiling is critical for understanding disease heterogeneity, progression and treatment resistance.

Single-cell DNA sequencing (scDNA-seq) enables genomic profiling of individual cells by directly measuring read depth at cellular resolution, thereby allowing the study of disease progression (Navin et al. 2011; Baslan and Hicks 2017). Numerous methods have been developed for CNA detection from scDNA-seq data, including HMMcopy (Shah et al. 2006), SCOPE (Wang et al. 2020), SCONCE (Hui and Nielsen 2022), SCICoNE (Kuipers et al. 2025) and rcCAE (Yu et al. 2023b). However, technical noise introduced by whole-genome amplification, such as GC-content bias and uneven coverage (Mallory et al. 2020), makes it difficult to distinguish true CNAs from amplification artifacts. Transcriptomic data also reflect copy number through gene dosage effects (Bhattacharya et al. 2020), which has motivated single-cell RNA sequencing (scRNA-seq) methods including CopyKAT (Gao et al. 2021) and SCEVAN (De Falco et al. 2023). However, scRNA-seq captures only expressed genes and produces sparse and locally dependent signals (Reyal et al. 2005). Moreover, RNA-seq measures gene expression rather than DNA copy number directly, which makes CNA inference highly dependent on transcriptional activity.

Emerging single-cell multimodal sequencing technologies, such as scTrio-seq2 (Bian et al. 2018) and scONE-seq (Yu et al. 2023a), have enabled simultaneous profiling of DNA and RNA within the same cell, providing a unique opportunity to jointly analyze complementary modalities for CNA inference. Integrative approaches have begun to explore this direction. For example, CONGAS+ (Patruno et al. 2023) uses a Bayesian mixture model to infer copy number clones from single-cell RNA and chromatin accessibility data (i.e., ATAC-seq). It models each omics layer with its own count likelihood and links the two modalities through shared latent copy number states. Numbat-multiome (Li et al. 2025a) aggregates single-cell RNA and ATAC-seq data into common genomic bins and infers allele-specific copy number using a joint hidden Markov model. However, both methods were developed primarily for RNA and ATAC-seq integration, and modality-specific differences in sparsity, coverage and measurement characteristics are not explicitly modeled within a unified framework.

Denoising sparse single-cell signals and effectively integrating multi-omics information are the other important tasks in CNA detection. Deep learning approaches, including autoencoder-based methods such as rcCAE (Yu et al. 2023b) and DeepCNA (Liu et al. 2024a) have improved signal quality, but their convolutional architectures do not naturally prioritize informative regions. In addition, their mean squared error (MSE)-based reconstruction losses penalize breakpoint transitions and stochastic noise equally, which often blurs sharp CNA boundaries. Moreover, existing integrative frameworks such as CONGAS+ and Numbat-multiome either use modality-specific likelihood weighting (Patruno et al. 2023) or aggregation of modality data for joint inference (Li et al. 2025a). However, modality-specific sparsity and uneven coverage, which are particularly present in scRNA-seq, are not explicitly modeled in these integration strategies.

Here, we present MintCNA, a unified framework for CNA profiling that integrates paired single-cell multi-omics data. MintCNA is novel in two aspects. First, it denoises signals through an attention-guided denoising convolutional autoencoder (AG-DCAE) with an edge-preserving loss function that avoids over-smoothing across breakpoint transitions while preserving the piecewise-constant CNA structure. Second, it applies a multivariate change-point procedure that jointly integrates signals across omics layers while accounting for structured missingness in omics layers. We conducted simulations and applied MintCNA to two human cancer datasets, and benchmarked it against existing methods in both unimodal (e.g., when RNA-seq was absent) and multimodal/multi-omics settings. As paired multi-omics sequencing technologies become increasingly accessible, MintCNA provides a tool for integrative CNA profiling to study disease progression and tumor evolution.

## Results

### Overview of the MintCNA framework

MintCNA is a unified CNA profiling framework for single-cell multi-omics data that combines deep-learning-based denoising and multivariate change-point detection (Fig. 1). MintCNA proceeds through four steps: (1) quality control and normalization of the input signal; (2) noise reduction by an attention-guided denoising convolutional autoencoder (AG-DCAE) that reduces technical noise while preserving CNA boundaries; (3) breakpoint detection through a multivariate change-point procedure that pools information across omics layers; and (4) CNA state assignment via a truncated Gaussian mixture model (TGMM). As a downstream step, subclones are identified by K-means clustering of the AG-DCAE latent representations.

**Figure 1.**
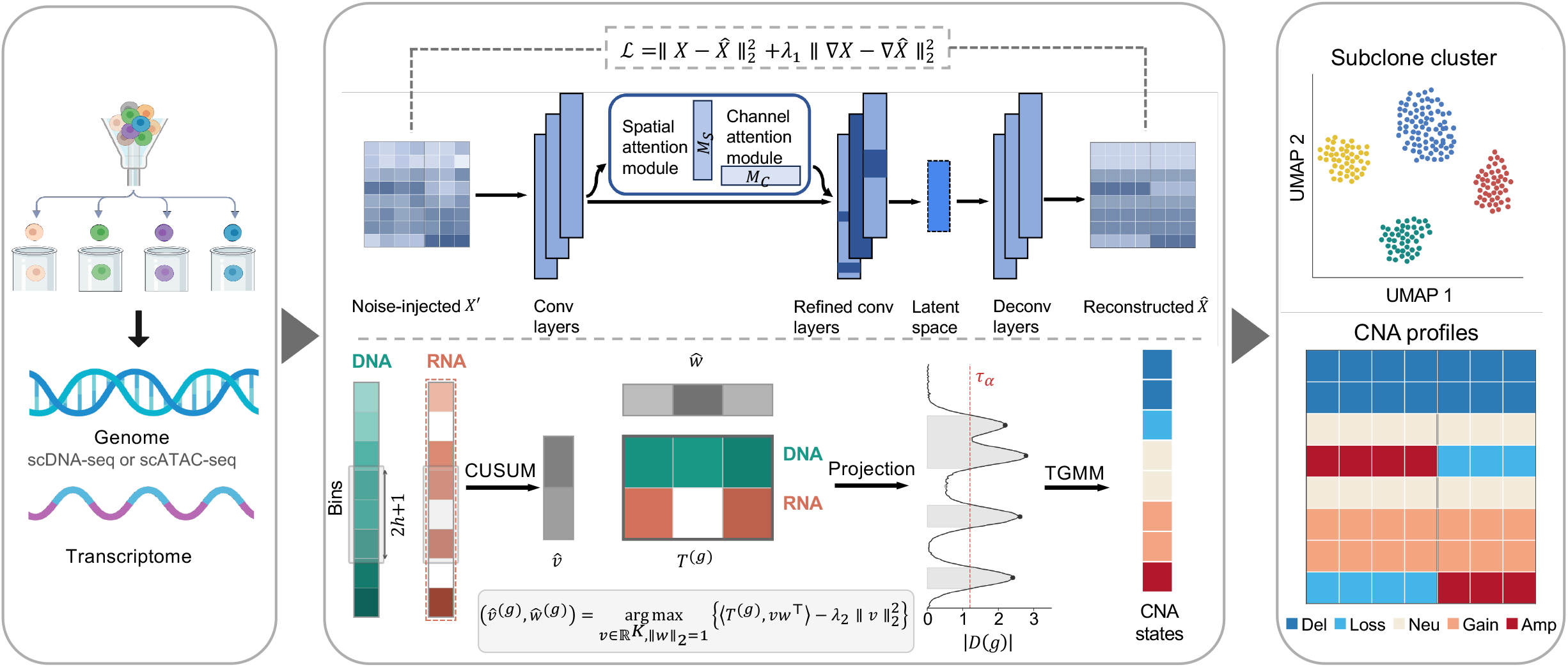
Overview of the MintCNA framework. MintCNA jointly models paired single-cell genomic and transcriptomic data for CNA detection. In the AG-DCAE denoising step (top), Gaussian noise is added to the *L2R* profiles during training. A convolutional autoencoder with a convolutional block attention module (CBAM) produces denoised *L2R* profiles. The edge-preserving loss function ℒ (dashed box) preserves sharp transitions at CNA boundaries. In the segmentation step (bottom), a sliding window scans the denoised genomic and transcriptomic data along the genome. The two signals are aggregated via a multivariate projection into a diagnostic score |D(g)|, and change-point candidates exceeding the threshold *τ*_*α*_ are considered breakpoints. Segments are classified by a truncated Gaussian mixture model (TGMM) into five CNA states, deletion (Del, CN = 0), loss (CN = 1), neutral (Neu, CN = 2), gain (CN = 3) and amplification (Amp, CN ≥ 4). Subclonal populations are identified by K-means clustering of the AG-DCAE latent representations.

### Denoising single-cell signals while preserving chromosomal breakpoint structure

Copy number signals are piecewise constant along the genome (Olshen et al. 2004), usually with sharp transitions at CNA boundaries that whole-genome amplification noise can obscure. Denoising is therefore essential to improve the signal-to-noise ratio without smoothing away the true chromosomal breakpoints. rcCAE (Yu et al. 2023b) addresses this with a convolutional autoencoder (CAE) that encodes the signal into a compact latent representation and reconstructs a denoised signal by minimizing mean squared error (MSE), 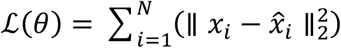. However, its loss function may blur the CNA breakpoints by generating an overly smooth reconstruction (Supplementary Fig. 1), and the CAE architecture weights all bins equally without positional selectivity. We therefore develop an attention-guided denoising convolutional autoencoder (AG-DCAE) whose loss function adds an edge-preserving penalty to the MSE, 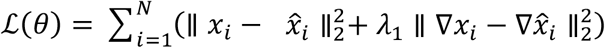. The second term penalizes any difference between the denoised and original profiles in the *L2R* differences between adjacent genomic bins. As a result, the model preserves abrupt copy number transitions instead of smoothing them away. Originally introduced to preserve sharp edges in predicted video frames (Mathieu et al. 2016) and later applied to preserve waveform shape in one-dimensional electrocardiogram signals (Liu et al. 2024b), this loss is well suited to retaining true CNA boundaries.

Furthermore, to allow the model to focus on informative genomic regions, we incorporate a convolutional block attention module (CBAM; Woo et al. 2018) into the convolutional encoder. CBAM sequentially applies channel attention and spatial attention. Channel attention uses maximum and average pooling across genomic bins to identify and emphasize informative feature maps, whereas spatial attention assigns weights across genomic bins to highlight regions likely to contain copy number transitions while suppressing noisy signals. As such, this dual-attention module is well suited for CNA denoising while introducing minimal computational burden. In our simulations, MintCNA showed denoised signals that more closely matched the ground truth than the MSE-based approach originally proposed in rcCAE (Yu et al. 2023b) and better preserved the true CNA boundaries (Supplementary Fig. 1), thereby improving breakpoint localization and CNA detection.

### Missingness-adjusted multivariate change-point detection integrating multi-omics datasets

The second main innovation of MintCNA is a multivariate change-point procedure that jointly detects CNA breakpoints across multi-omics profiles. A challenge in omics integration is the heterogeneity across omics layers. For example, RNA-seq provides coverage only for expressed genes, resulting in substantial missingness across genomic bins. Moreover, integrating complementary signals from different modalities requires a unified statistic that captures shared signals of copy number changes. To address these challenges, we develop a multivariate change-point detection approach that accommodates heterogeneous missingness in high-dimensional data (Follain et al. 2022).

A CNA breakpoint corresponds to a genomic position at which the mean signal differs between the flanking chromosomal segments. MintCNA detects such changes using a cumulative sum (CUSUM) statistic, which contrasts the mean signals on the two sides of a candidate position. To identify multiple breakpoints across the genome, we compute the statistic within sliding genomic windows and select local maxima as candidate breakpoints. We further adjust the CUSUM statistic for layer-specific missingness by accounting for the number of observed bins on either side of a candidate position, thereby mitigating the influence of uneven genomic coverage.

To integrate signals across omics layers, MintCNA combines the layer-specific CUSUM statistics into a unified diagnostic score using an *l*_2_-regularized rank-1 projection. The projection weights are estimated adaptively from the data, so that each omic layer contributes according to the strength of its breakpoint signals. Candidate positions with diagnostic scores exceeding a threshold derived from an empirical null distribution are identified as breakpoints (Methods). Together, the missingness-adjusted CUSUM statistic and regularized multivariate projection enable MintCNA to integrate complementary signals across omics layers while accounting for differences in genomic coverage.

### MintCNA detected CNAs most accurately across simulation scenarios

We first evaluated MintCNA against existing scDNA-seq CNA callers using simulated datasets. MintCNA achieved the highest overall accuracy for both deletions and duplications across the evaluated scenarios (Fig. 2, Supplementary Tables 1–3). For deletions, MintCNA achieved an F1 score of 0.76 under the most favorable scenario (0.2× coverage, no additional noise) and 0.65 under the most challenging scenario (0.05× coverage, noise level 0.1). rcCAE as another autoencoder-based denoising method was the next most accurate one for deletions (F1 = 0.49– 0.64). HMMcopy, SCOPE, and SCCNV showed high recall (0.63–0.96) but low precision (0.08– 0.24), indicating a high proportion of false-positive calls (Supplementary Tables 1–3). SCICoNE detected relatively few events (recall = 0.08–0.19) and had consistently low F1 scores (0.12–0.24), whereas SCONCE showed variable performance across scenarios (F1 = 0.18–0.45).

**Figure 2.**
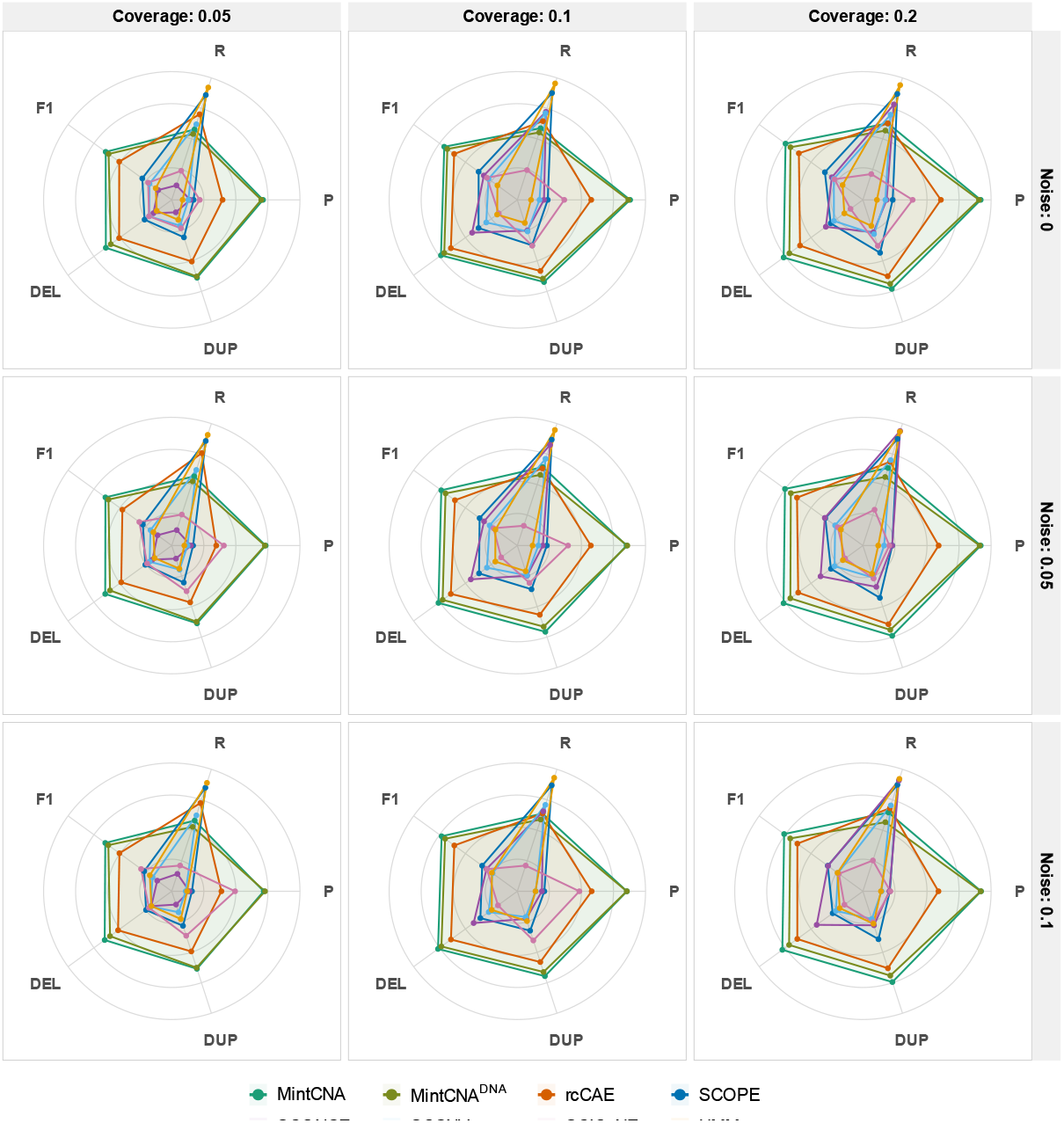
Event-level detection accuracy across simulation scenarios. Columns represent sequencing coverage (0.05×, 0.1×, and 0.2×) and rows represent noise levels (0, 0.05, and 0.1). Radar plots show overall precision (P), recall (R), and F1, together with F1 scores for deletions (DEL) and duplications (DUP). Concentric rings mark values of 0.25, 0.50, 0.75 and 1.00.

Stratification analyses by CNA size showed that the relative advantage of MintCNA was greater for smaller events (Supplementary Figs. 2–4). MintCNA, rcCAE, and SCOPE detected large CNAs (>10 Mb) with comparable accuracy, whereas MintCNA achieved the highest accuracy for medium (3–10 Mb) and focal (<3 Mb) events, for which breakpoints are more difficult to distinguish from technical noise. Despite its higher event-level detection accuracy, MintCNA maintained breakpoint localization comparable to the other leading methods. Mean boundary bias ranged from 2.5 to 3.6 Mb for MintCNA and from 2.1 to 3.0 Mb for rcCAE across simulation scenarios (Fig. 3a). MintCNA also detected copy number profiles most closely consistent with the ground truth, with mean root mean squared error (RMSE) ranging from 0.18 to 0.23, compared with 0.21–0.28 for rcCAE and 0.31–0.75 for the remaining methods (Fig. 3b).

**Figure 3.**
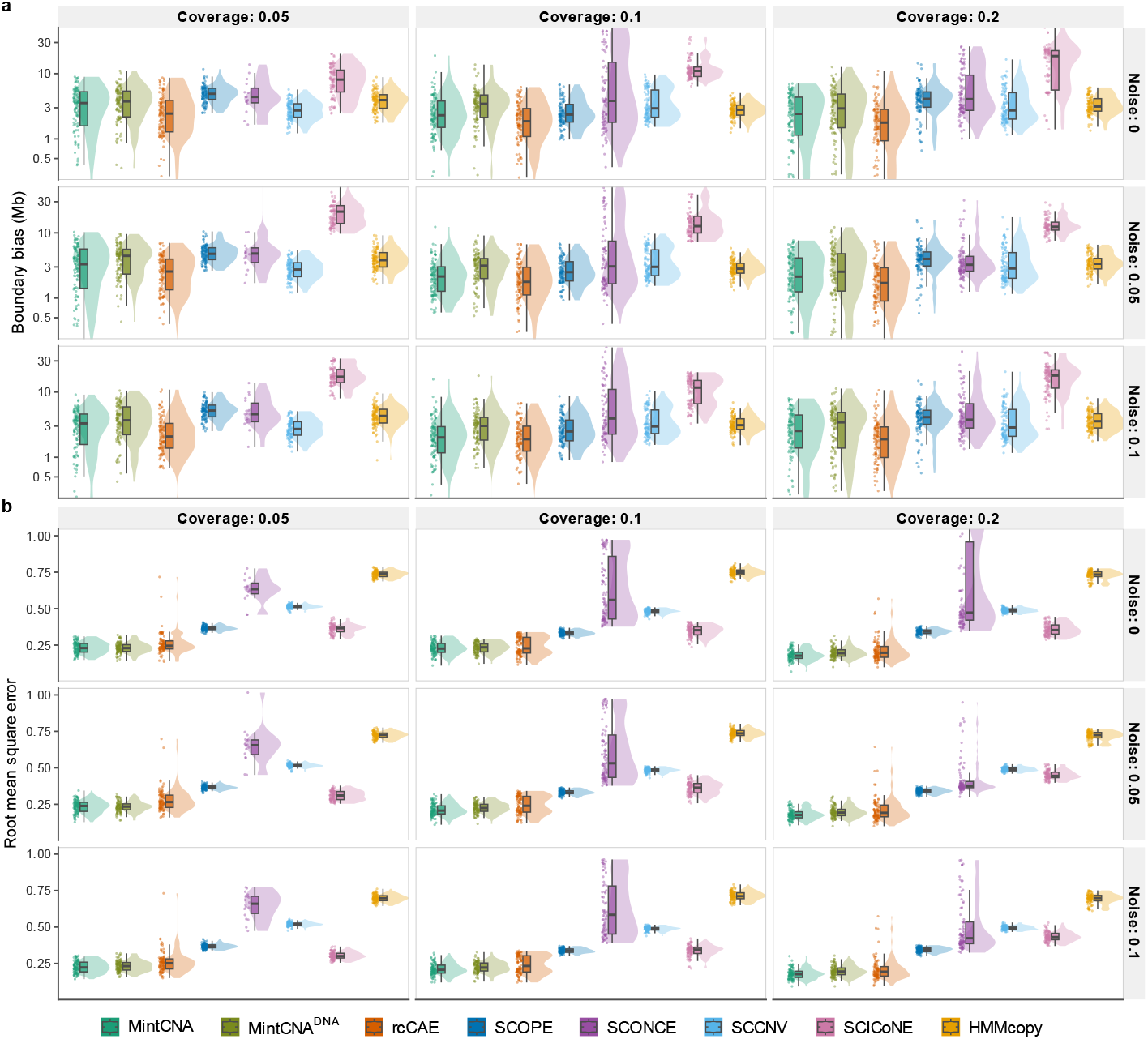
Breakpoint localization and copy number accuracy across simulation scenarios. **(a)** Boundary bias (Mb), averaged across true positive CNA events within each cell and shown on a log scale. **(b)** Bin-level root mean squared error (RMSE) between predicted and ground-truth copy number states. Metrics are shown across three coverage depths (0.05×, 0.1×, and 0.2×) and three noise levels (0, 0.05, and 0.1) for all evaluated methods. Each point represents one of the 90 tumor cells in each scenario, and only cells with true-positive events are shown in (a).

To assess the contribution of multi-omics integration, we additionally evaluated a DNA-only mode that uses scDNA-seq data alone, denoted by MintCNA^DNA^. MintCNA^DNA^ achieved F1 scores of 0.59–0.73 for deletions and 0.62–0.69 for duplications and remained more accurate than the other scDNA-seq callers across all evaluated scenarios (Supplementary Tables 1 and 2). Adding scRNA-seq data increased F1 scores by 0.01–0.07 relative to MintCNA^DNA^. In contrast, the RNA-only mode, MintCNA^RNA^, showed substantially lower precision and recall than either the integrated or DNA-only modes across all scenarios (Supplementary Fig. 5b–d). Together, these results indicate that the DNA layer provides the primary signal for CNA detection, while integration of the RNA signal provides additional information that modestly improves detection accuracy.

### MintCNA revealed progressive CNA evolution in a colorectal cancer multi-omics dataset

Colorectal cancer (CRC) is the second-leading cause of cancer-related death worldwide, with metastasis accounting for most CRC-related mortality (Siegel et al. 2023). CNAs contribute substantially to intratumor heterogeneity (ITH) and have been implicated in metastatic progression and therapeutic resistance (Mamlouk et al. 2017; The Cancer Genome Atlas Network 2012). To characterize the CNA evolution in metastatic CRC at single-cell resolution, we applied MintCNA to scTrio-seq2 data from patient CRC01, which assessed paired DNA and RNA profiles of 196 cells across four tumor sites (Bian et al. 2018). The four sites were the primary tumor (PT), lymph node metastasis (LN), liver metastasis (ML), and post-treatment liver metastasis (MP), with 59, 7, 55 and 75 cells, respectively.

MintCNA resolved the CNA landscape of CRC01 into clonal alterations shared across tumor sites and subclonal alterations that varied among sites (Fig. 4a). Clonal losses involved 8p and chromosomes 15, 18 and 22, including loss of the tumor suppressor genes *SMAD4* and *DCC* at 18q21 (The Cancer Genome Atlas Network 2012). Recurrent gains on 8q, chromosome 13 and 20q were observed across all four sites, including amplification of *MYC* at 8q24 and gains involving the CRC-associated genes *CDX2* (13q12), *AURKA* (20q13), and *SRC* (20q11) (Salari et al. 2012; Sheffer et al. 2009). Together, these recurrent alterations were consistent with the chromosomal instability landscape characteristic of colorectal tumors.

**Figure 4.**
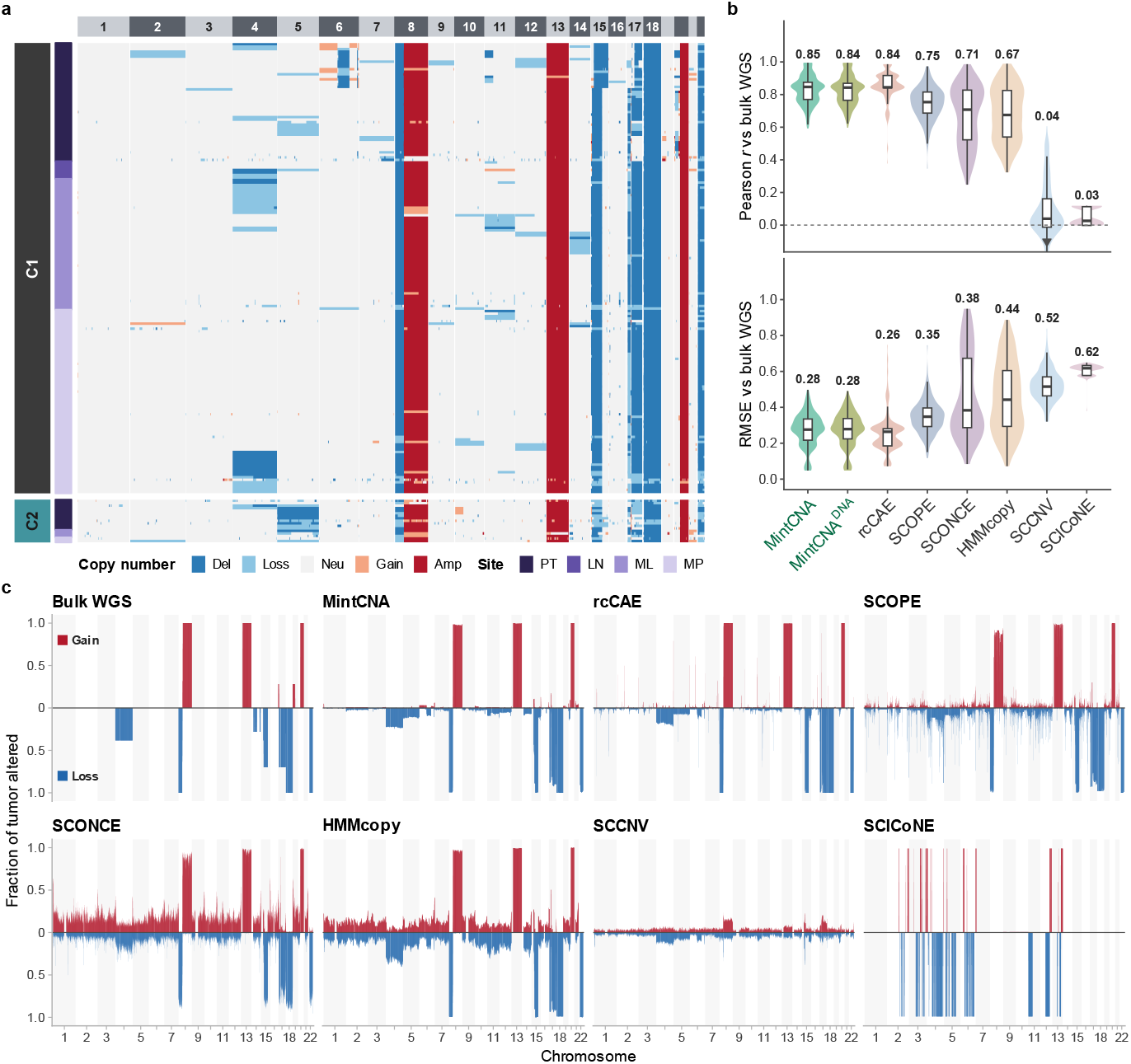
Application of MintCNA to the colorectal cancer scTrio-seq2 dataset. **(a)** Genome-wide CNA profiles inferred by MintCNA for 196 cells from patient CRC01, ordered by subclone assignment (C1 and C2) and tumor site: primary tumor (PT), lymph node metastasis (LN), liver metastasis (ML), and post-treatment liver metastasis (MP). Copy number states are deletion (copy number [CN]=0), loss (CN=1), neutral (CN=2), gain (CN=3), and amplification (CN≥4). **(b)** Pearson correlation (upper) and root mean squared error (RMSE; lower) between the single-cell CNA profile inferred by each method and the matched bulk WGS profile from the corresponding tumor site (n = 196 cells). **(c)** Pseudobulk CNA profiles inferred by each method compared with the matched bulk WGS reference. For each genomic bin, the fractions of tumor cells called as having a gain (red, above) or loss (blue, below) are shown. For bulk WGS, each site-level profile is weighted by the number of cells from that site.

Beyond these shared alterations, chromosome 17 loss represented a major source of ITH and showed a marked increase in prevalence from the primary tumor to the metastatic sites (Supplementary Fig. 6). In the primary tumor, 66.1% and 69.5% of cells carried 17p and 17q losses, respectively, compared with 85.7%–100% across the metastatic sites. K-means clustering of the AG-DCAE latent representations identified two subclones (Fig. 4a; Methods). The dominant subclone C1 was present across all tumor sites and carried 17q loss, whereas C2 was largely restricted to the primary tumor and additionally carried 5q loss encompassing the WNT pathway regulator *APC*. The enrichment of the chromosome 17-loss lineage in the metastatic sites, together with the restriction of C2 largely to the primary tumor, is consistent with preferential dissemination or expansion of C1 during metastasis (Pino and Chung 2010).

To evaluate CNA detection performance, we compared the single-cell CNA profiles inferred by each method with the matched bulk WGS profiles of the corresponding tumor sites (Fig. 4b). MintCNA and rcCAE showed the highest concordance with bulk WGS, with median Pearson correlations of 0.85 and 0.84 and median RMSEs of 0.28 and 0.26, respectively. MintCNA^DNA^ performed comparably to the full MintCNA model (median r = 0.84), likely because the CNA landscape of the patient was dominated by arm-level and whole-chromosome alterations that can be reliably captured using the DNA signal alone. SCOPE, SCONCE, and HMMcopy showed lower concordance (r = 0.67 – 0.75) (Fig. 4b) and inferred numerous gains and losses that were not indicated by bulk WGS (Fig. 4c). SCCNV (r = 0.04) detected few of the clonal CNAs, whereas many losses inferred by SCICoNE (r = 0.03) were absent from the bulk WGS profiles. These results support the accuracy of MintCNA in recovering the major CNA landscape observed in bulk WGS.

### MintCNA identified copy number subclones associated with distinct malignant cell states in IDH-mutant glioma

The design of MintCNA enables CNA detection in single-cell modalities beyond paired scDNA-seq and scRNA-seq. We further applied MintCNA to 10x Multiome data from the CARE IDH-mutant glioma cohort with paired scATAC- and scRNA-seq data (Johnson et al. 2026). In IDH-mutant gliomas, CNAs define two major molecular subtypes, as the 1p/19q codeletion distinguished oligodendroglioma and astrocytoma (Louis et al. 2021). The analyzed cohort comprised 47,210 malignant nuclei from 20 samples of 10 patients after quality control (Supplementary Fig. 7).

We compared MintCNA and its single-omic mode MintCNA^ATAC^ with existing single-omics methods (i.e., for ATAC-seq or RNA-seq) and multi-omics methods (i.e., CONGAS+) using matched bulk WGS profiles as a reference. MintCNA achieved the highest median Pearson correlation with bulk WGS (r = 0.80; Fig. 5a) and the highest F1 score (0.59; Fig. 5b). MintCNA^ATAC^ performed comparably to the full model (r = 0.78), likely because the CNA landscape in this cohort is dominated by arm-level events that can be captured from ATAC fragment coverage alone. The ATAC-seq method AtaCNA achieved a similar correlation (r = 0.77) and higher precision (0.83) but had lower recall than MintCNA (0.39 versus 0.51). Copy-scAT (r = 0.68) called 19q loss in four of the six samples for which bulk WGS showed no deletion. RIDDLER (r = 0.61) had limited recall (0.24), whereas epiAneufinder (r = 0.57) and the RNA-based caller inferCNV (r = 0.33) had recall below 0.01. The omics integration method CONGAS+ (r = 0.11) detected no copy number losses using the package’s arm-level segmentation settings. Collectively, these results demonstrate that MintCNA generalizes to chromatin accessibility and transcriptomic data without modification of the overall framework.

**Figure 5.**
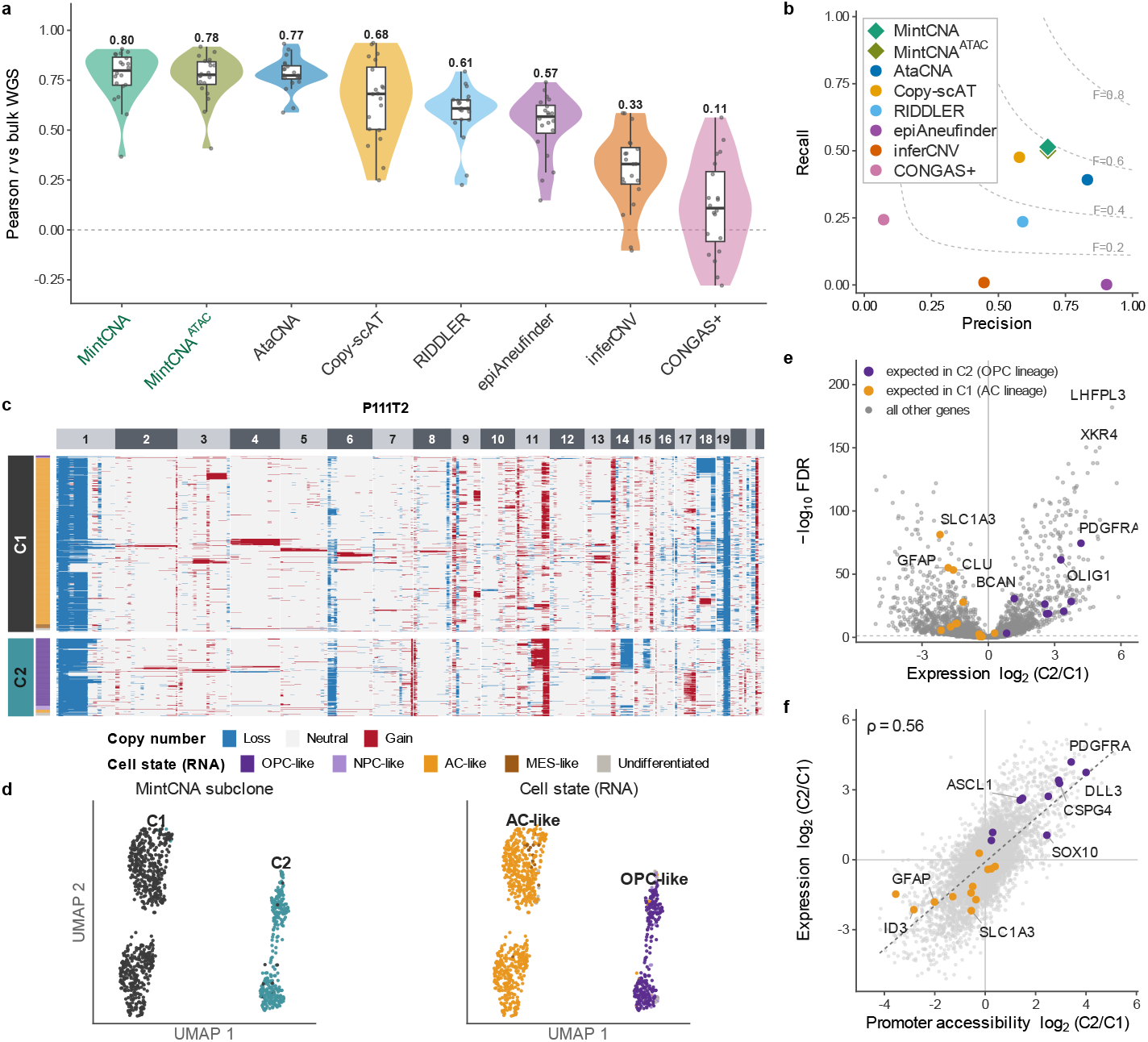
Application of MintCNA to the CARE IDH-mutant glioma 10x Multiome dataset. **(a)** Pearson correlation between the pseudobulk copy number profile inferred by each method and the matched bulk WGS profile across 20 samples. Each point represents one tumor sample. **(b)** Bin-level precision and recall against matched bulk WGS, pooled across 20 samples. Dashed curves indicate contours of constant F1 score. **(c)** Genome-wide copy number profiles inferred by MintCNA for 995 malignant nuclei from sample P111T2, ordered by subclone assignment. Copy number states are shown as loss (CN=1), neutral (CN=2) and gain (CN=3). Cell state annotations from the source study are shown alongside MintCNA subclone assignments. **(d)** RNA UMAP of the same nuclei, colored by MintCNA subclone assignment (left) and cell-state annotation from the source study (right). **(e)** Differential expression between subclones C2 and C1 using the Wilcoxon rank-sum test with Benjamini–Hochberg correction. The FDR threshold was 0.05. **(f)** Correlation between change in expression and change in promoter accessibility between the two subclones. ρ, Spearman correlation. In (e) and (f), lineage genes are colored by the subclone in which they are expected to be more highly expressed.

Beyond CNA detection accuracy, we examined whether the subclones detected by MintCNA were consistent with the RNA-derived malignant cell states reported in the source study (Johnson et al. 2026). In sample P111T2, an oligodendroglioma comprising 995 malignant nuclei, MintCNA detected the hallmark 1p/19q codeletion in nearly 90% of nuclei in both subclones (Fig. 5c). The two subclones identified from the AG-DCAE latent representation showed strong consistency with the originally reported cell states, with 95.0% of C1 nuclei classified as astrocyte-like (AC-like), and 87.1% of C2 nuclei classified as oligodendrocyte progenitor-like (OPC-like). In RNA expression space (Fig. 5d), a nearest-neighbor classifier predicted the MintCNA subclone labels (i.e., C1 or C2) with 98.3% accuracy, compared with 99.4% for the cell state annotations (i.e., AC-like or OPC-like) defined from the same data. Thus, subclonal structure identified independently from ATAC-derived copy number profiles closely recapitulated transcriptionally defined malignant cell states.

We next characterized the transcriptional and epigenomic differences between the two subclones. Of the 10,756 genes tested, 4,289 were differentially expressed (FDR < 0.05; Fig. 5e), with the oligodendroglial markers *PDGFRA, OLIG1* and *BCAN* (Neftel et al. 2019) among the strongly upregulated genes in C2. Between the two subclones, differences in promoter accessibility were significantly correlated with differences in gene expression (Spearman ρ = 0.56; Fig. 5f). *SOX10* and *SOX8* motifs were more accessible in C2, whereas five *NFI* motifs were more accessible in C1 (Supplementary Fig. 8a), consistent with the known roles of *NFI* in astrocyte specification and *SOX10* in oligodendrocyte specification (Glasgow et al. 2014). These differences were not explained by gene dosage, as 15 lineage genes had the same copy number in both subclones, and the correlation between promoter accessibility and gene expression remained unchanged after excluding genes with different copy number (Supplementary Fig. 8b, c). These findings indicate that the MintCNA-defined subclones capture distinct oligodendroglial and astrocytic regulatory states beyond differences attributable to copy number alone.

### Computational efficiency comparison

We compared the computational efficiency of MintCNA with that of the benchmarked methods using the simulated, colorectal and glioma datasets. All methods received the same input read-count matrix where applicable, and runtime was measured from input processing to generation of the final CNA profiles. Analyses were performed on a high-performance computing cluster using eight CPU cores and 64 GB of RAM. MintCNA analyzed the simulated and colorectal cancer datasets in 28 and 80 min, respectively. It was at least four-fold faster than SCOPE, SCONCE, and SCCNV, but slower than rcCAE which required 25 and 45 min, respectively (Supplementary Fig. 9a). HMMcopy and SCICoNE were the fastest methods (4–14 min) but showed lower CNA detection accuracy in our simulations.

On the glioma 10x Multiome cohort, MintCNA required a median of 145 CPU core-minutes per tumor, compared with 1.7–16.5 for the other methods and 241 for epiAneufinder (Supplementary Fig. 9b). The greater computational cost of MintCNA in this setting was primarily attributable to its tumor-specific AG-DCAE training procedure and change-point scanning across individual nuclei. Overall, MintCNA showed a moderate computational cost, with runtime varying according to dataset size and modality.

## Discussion

In this study, we developed MintCNA, a unified framework for CNA detection from paired single-cell multi-omics data. MintCNA is the first computational tool that utilizes the advantages of statistical modeling and deep learning in efficient CNA detection with paired single-cell omics data. It makes two main methodological contributions by jointly modeling paired molecular layers from the same cell and incorporating an attention-guided denoising procedure to enhance CNA signals in the preprocessing steps. Comprehensive simulations showed MintCNA’s superior detection accuracy across various scenarios. Application to a scTrio-seq2 colorectal cancer dataset with paired single-cell DNA and RNA level data, and an IDH-mutant glioma cohort with paired single-cell ATAC-seq and RNA-seq data, showed its generalizability across different omics types.

Beyond accurate CNA detection, we showed that CNAs were associated with metastasis in colorectal cancer and subclones were linked to distinct cell states in glioma. In CRC01, chromosome 17 loss was present in most metastatic cells but in fewer primary tumor cells. This indicates that most metastatic cells derived from one lineage with chromosome 17 loss, which covers *TP53* on 17p and is common in colorectal cancer (Pino and Chung 2010). Single-cell CNA profiles across tumor sites can thus help trace the lineages from which metastases arise. In glioma, the two subclones identified from the latent representation corresponded to the AC-like and OPC-like states. Like rcCAE (Yu et al. 2023b), MintCNA infers subclones directly from read coverage so the subclones can capture both genetic and regulatory heterogeneity between cells. The subclones may therefore help elucidate genetic evolution across cell-state changes during glioma progression (Johnson et al. 2026).

Technical noise arising from whole-genome amplification and uneven sequencing coverage remains a major challenge for single-cell CNA detection. Existing methods address this challenge through different strategies. Ginkgo (Garvin et al. 2015) corrects for GC-content bias, whereas SCOPE (Wang et al. 2020) accounts for technical artifacts using a latent-factor model anchored on normal control cells. More recently, rcCAE (Yu et al. 2023b) introduced a convolutional autoencoder to denoise single-cell read counts. MintCNA introduces an attention-guided denoising convolutional autoencoder (AG-DCAE) to suppress technical noise while retaining abrupt transitions at CNA boundaries. CBAM (Woo et al. 2018) further enables adaptive weighting of learned features and genomic positions, so that the model focuses on informative signals. To our knowledge, MintCNA is the first framework to combine attention-guided denoising with an edge-preserving loss for single-cell CNA detection. Together, these components reduce technical noise while preserving the genomic structure relevant to CNA segmentation.

Although each molecular layer (e.g., DNA, RNA, chromatin accessibility) provides information about underlying copy number alterations, single-modality methods may not fully exploit the comprehensive signals available from multi-omics. MintCNA addresses this limitation by combining signals across omics layers into a unified breakpoint score which allows signals from the two modalities to complement each other. Simulations showed consistently improved CNA detection performance of MintCNA over single-omic approaches in all scenarios. In both real-world datasets, MintCNA achieved the highest concordance with matched bulk WGS, although performance was comparable to the corresponding single-omic model in datasets dominated by broad chromosomal alterations. These findings suggest that multi-omics integration may be particularly beneficial under limited sequencing coverage or for detecting focal CNAs, where complementary information from a second modality can strengthen breakpoint signals.

One limitation of the current framework is that MintCNA performs cell-specific CNA segmentation, which does not explicitly use shared CNA patterns across cells from the same clone. Previous methods, including SeCNV (Wang et al. 2022) and HiScanner (Zhao et al. 2025), have shown the potential benefits of pooling information across cells to improve sensitivity to clonal events. Given that the AG-DCAE latent representations can capture subclonal structure, a promising extension would be to identify putative subclones from the latent representations before jointly segmenting cells within each group. This could improve the detection of shared breakpoints across cells while retaining the ability to characterize intratumor CNA heterogeneity. Furthermore, the current latent representations are learned from the coverage-based layer alone. Incorporating a separate latent space for the RNA layer could make subclone clustering more comprehensive.

In addition, the successful application of MintCNA to 10x Multiome data shows the generalizability of the framework across different combinations of molecular modalities and suggests its potential applicability to other paired ATAC and RNA profiling technologies, including SHARE-seq (Ma et al. 2020) and SNARE-seq (Chen et al. 2019). With the emergence of high-throughput paired DNA and RNA co-sequencing technologies, such as DEFND-seq (Olsen et al. 2025) and BAG-seq (Li et al. 2025b), MintCNA offers a computational approach for leveraging these increasingly accessible multi-omics datasets. These advances may facilitate more comprehensive characterization of CNA heterogeneity and its role in malignancy across diverse tumor types.

## Methods

### Data preprocessing

MintCNA is designed for paired scDNA-seq and scRNA-seq data and extends naturally to other multi-omics settings that combine a coverage-based genomic layer with a transcriptomic layer (e.g., 10x Multiome scATAC-seq and scRNA-seq).

For the coverage-based genomic data (i.e., scDNA-seq), reads are counted in fixed-width, non-overlapping genomic bins (e.g., 100 kb). Bins with extreme GC content or low mappability are excluded, and the remaining read counts are corrected for GC-content and mappability biases using a median-normalization procedure (Magi et al. 2013). The adjusted counts are then normalized to counts per million (CPM) and transformed into log2 ratios (*L2R*) relative to the median across normal cells within each bin. When annotated normal cells are unavailable, putative diploid cells are identified computationally based on coverage uniformity (Wang et al. 2020).

For the transcriptomic data (i.e., scRNA-seq), we retain cells with > 200 detected genes and genes detected in > 5% of cells. Following CopyKAT (Gao et al. 2021), each expression profile is variance-stabilized using the log–Freeman–Tukey transformation and smoothed along genomic coordinates using a first-order polynomial dynamic linear model. Gene-level *L2R* values are then calculated relative to the median expression across normal cells. To enable joint modeling of the two omics layers, the genomic and transcriptomic signals are mapped to a common set of genomic bins. Gene-level *L2R* values are assigned to every genomic bin overlapping the corresponding gene, and multiple genes within a bin are aggregated by their median. Bins without overlapping genes are treated as missing in the RNA layer.

### Data notations and models

Let *X*_*i*_ ∈ ℝ^*K*×*G*^ denote the *L2R* matrix of the *i*-th cell (*i* = 1, …, *N*) after preprocessing, where *N* is the number of cells, *K* is the number of omics layers and *G* is the total number of genomic bins. We model each omics sequence along the genome as

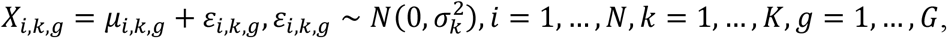

where *μ*_*i,k,g*_ is the underlying piecewise-constant mean along the genome and 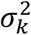 denotes the noise variance of omics layer *k*. A change-point is a position *g*\* where 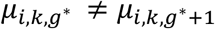. Because coverage and dropout rates differ across omics layers, some bins of *X*_*i*_ are unobserved. We encode this missingness pattern with a binary observation indicator Ω_*i*_ ∈ {0,1}^*K*×*G*^, where Ω_*i,k,g*_ = 1 when omics layer *k* is observed at bin *g*, and Ω_*i,k,g*_ = 0 otherwise.

### Attention-guided denoising convolutional autoencoder for CNA signal enhancement

To denoise the dataset for enhanced CNA signal, we introduce an attention-guided denoising convolutional autoencoder (AG-DCAE) for the coverage-based *L2R* profiles. We train the network as a denoising autoencoder by adding Gaussian noise to the input *L2R* profile. Specifically, for cell *i*, let *x*_*i*_ = (*X*_*i*,1,1_, …, *X*_*i*,1,*G*_)^⊤^ denote its coverage-based *L2R* profile, and a noisy profile is given by *x*′ = *x*_*i*_ +*η*_*i*_ with 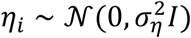.

We use the encoder *f*_*θ*_(⋅) to map the noisy profile 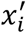 to a low-dimensional latent representation 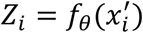 using stacked one-dimensional convolutional layers followed by a convolutional block attention module (Woo et al. 2018). CBAM is applied to the feature map of the final convolutional layer, before the fully connected bottleneck that produces *Z*_*i*_. It sequentially applies channel attention which weights the learned feature maps based on maximum- and average-pooled features across genomic bins, and spatial attention which assigns weights across genomic positions based on local feature information. *Z*_*i*_ is then passed through a symmetric transposed-convolutional decoder *d*_*θ*_(⋅) to reconstruct the denoised *L2R* profile 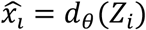.

The training objective combines a reconstruction loss between the original and reconstructed profiles with an edge-preserving penalty (Mathieu et al. 2016):

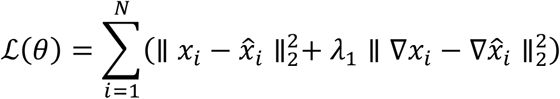

where ∇*x*_*i*_ denotes the gradients between adjacent genomic bins and *λ*_1_ controls the weight of the edge-preserving term. The second term penalizes discrepancies in adjacent-bin gradients between the original and reconstructed profiles, thereby preserving sharp CNA transitions during reconstruction. *θ* denotes the parameters of the convolutional encoder, the CBAM, and the decoder, which are jointly optimized by minimizing ℒ(*θ*). After training, the latent representation *Z*_*i*_ and the denoised *L2R* profiles 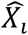 of each cell are obtained for subclone clustering and CNA detection. Detailed hyperparameter settings and the clustering procedure are provided in Supplementary Methods.

### Local CUSUM statistics with missingness adjustment

To detect breakpoints, we compute local CUSUM statistics along the genome from the denoised *L2R* matrix 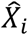 of each cell separately. Let *X*_*k,j*_ denote the denoised *L2R* signal of a cell for omics layer *k* at genomic bin *j*, and let Ω_*k,j*_ indicate whether the corresponding signal is observed. For each candidate breakpoint position *g*, we define a local window {*g* − *h*, …, *g* +*h*} centered at *g*. Within each window, we compute a missingness-adjusted CUSUM statistic for each omics layer *k*, following Follain et al. (2022) as

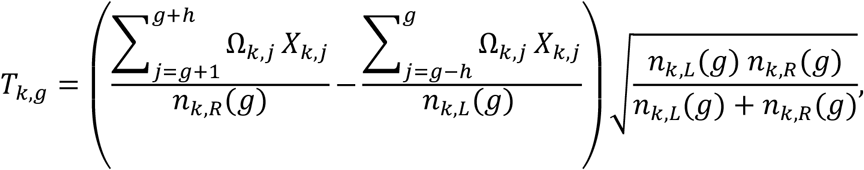

where 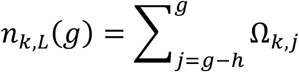 and 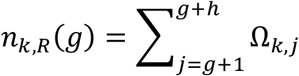 denote the numbers of observed bins to the left and right of *g*, respectively. The two fractions are the mean *L2R* values of the observed bins on either side of the candidate position, while the scaling factor 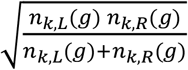 downweights the statistic at positions where one or both sides have sparse coverage.

### Multivariate projection for pooling cross-omics evidence

After the local CUSUM statistic computation, we integrate the statistics across omics layers to detect breakpoints. For each local window centered at genomic position *g*, we stack the local CUSUM statistics {*T*_*k,j*_} across the *K* omics layers and genomic bins within the window to form the matrix *T*^(*g*)^ ∈ ℝ^*K*×(2*h*+1)^. To pool the breakpoint signals from all omics layers into a unified score at each position, we approximate *T*^(*g*)^ ≈ *vw*^⊤^ through an *l*_2_-regularized rank-1 approximation (Follain et al. 2022) by solving

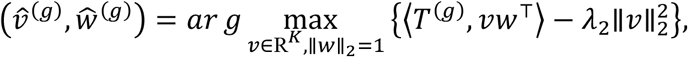

where 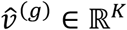 is an omics-weight vector that adaptively borrows information across omics layers, and *ŵ*^(*g*)^ ∈ ℝ^2*h*+1^ weights the bins within the window. Here the number of omics layers *K* is small and all layers are to be integrated. We therefore use an *l*_2_ (ridge) penalty 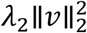 which regularizes the projection weights without explicitly inducing sparsity across omics layers. This optimization is biconvex and is solved by alternating closed-form updates of *v* and *w* until convergence. The integrated diagnostic score at position *g* is then defined as

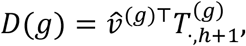

where 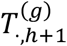 is the column of *T*^(*g*)^ at the center of the window. Repeating this procedure across genomic positions yields a genome-wide diagnostic sequence {*D*(*g*)}. A position *g* is identified as a breakpoint if

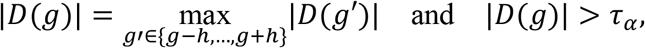

where *τ*_*α*_ is the (1-*α*) quantile of the empirical null distribution of the diagnostic sequence. The derivation of the empirical threshold, the closed-form alternating solution for the projection vectors and the choice of *λ*_2_ are provided in Supplementary Methods.

When only a single omics layer is available (*K* = 1), the multivariate projection is unnecessary as the local screening statistic *T*^(*g*)^ reduces to a vector and the diagnostic score reduces to *D*(*g*) = *T*_1,*g*_. The remaining steps of local maxima identification and significance thresholding proceed identically to the multi-omics setting.

### CNA calling via truncated Gaussian mixture modeling

After breakpoint detection, we assign each segment between adjacent breakpoints to a discrete CNA state using a truncated Gaussian mixture model (TGMM). In the multi-omics setting, TGMM is applied to the fused mean *Y*_*i,s*_ of segment *s* in cell *i*, which is the average of the layer-specific segment means weighted by the number of observed bins in each layer (Supplementary Methods). With a single omics layer, *Y*_*i,s*_ reduces to the segment mean of that layer.

Let *z*_*i,s*_ ∈ {1, …,5} denote the latent CNA state of segment *s* in cell *i*, corresponding to deletion (copy number [CN] = 0), loss (CN=1), neutral (CN=2), gain (CN=3), and amplification (CN ≥ 4) (Magi et al. 2013; De Falco et al. 2023). We model

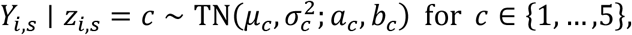

where TN(*μ*_*c*_, 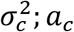, *b*_*c*_) denotes a truncated normal distribution with state-specific mean *μ*_*c*_, variance 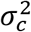 and truncation interval [*a*_*c*_, *b*_*c*_]. Model parameters are estimated using the expectation-maximization (EM) algorithm (Dempster et al. 1977), and each segment is assigned to the CNA state with the highest posterior probability. Adjacent segments assigned to the same state are subsequently merged to form the final CNA calls. Details of the EM algorithm and truncation intervals are provided in Supplementary Methods.

### Simulation design

We benchmarked MintCNA against existing CNA callers using simulated paired scDNA-seq and scRNA-seq data. Ground-truth single-cell CNA profiles were generated using CNAsim (Weiner and Bansal 2023), which simulates a cell-lineage tree with divergent subclones and CNA evolution along lineage branches. Read counts were simulated as RC_*i,g*_ ~ Poisson(κ ⋅ CN_*i,g*_ ⋅ *ω*_*g*_), where CN_*i,g*_ denotes the copy number of cell *i* at bin *g*, κ is the coverage scale factor, and *ω*_*g*_ models the uneven coverage introduced by whole-genome amplification using a Lorenz curve. The hg38 reference genome was used to define genomic bins and map genes to CNA regions. The baseline setting comprised 100 cells (10% normal), 100 kb bins, 0.1× coverage, three subclones, and no additional read count noise. To evaluate the effects of sequencing depth and technical noise on detection accuracy, we considered all combinations of three coverage depths (0.05×, 0.1×, and 0.2×) and three noise levels (0, 0.05, and 0.1). Noise was introduced by multiplying the read count of each bin by a Gaussian random factor with standard deviation equal to the noise level. Additional CNAsim parameters, including CNA lengths and event rates, are provided in Supplementary Methods.

Because existing simulators do not generate paired scDNA-seq and scRNA-seq data from the same cells, we simulated scRNA-seq counts for the corresponding cells using Splatter (Zappia et al. 2017). Baseline gene expression was first simulated and then modified according to the underlying copy number to introduce gene-dosage effects (Patruno et al. 2023). For a gene located within a segment with copy number *CN*, its baseline expression was multiplied by a random factor drawn from *N*(*CN*/2, 0.1^2^). Because not all genes exhibit expression changes proportional to copy number (Bhattacharya et al. 2020), we introduced a concordance indicator *B*_*i,m*_ ~ Bernoulli(*p*) for gene *m* in cell *i*, where *p* ∈ {0.25,0.5,0.75,1} specifies the probability that gene expression reflects the underlying copy number. For concordant genes (*B*_*i,m*_ = 1), the CNA-scaled expression was used, whereas baseline expression was retained for non-concordant genes (*B*_*i,m*_ = 0).

### Benchmarked methods and evaluation metrics

We compared MintCNA with six scDNA-seq CNA callers: rcCAE (Yu et al. 2023b), SCOPE (Wang et al. 2020), SCONCE (Hui and Nielsen 2022), SCCNV (Dong et al. 2020), SCICoNE (Kuipers et al. 2025) and HMMcopy (Shah et al. 2006). All benchmarked methods were run using their default settings unless otherwise specified (Supplementary Methods). Because the simulations were generated from DNA derived ground-truth CNA profiles, the primary benchmark focused on scDNA-seq CNA callers. For consistent evaluation across methods, inferred copy numbers were grouped into three states: deletion (CN < 2), neutral (CN = 2), and duplication (CN > 2).

We evaluated CNA detection using event-level precision, recall, and F1 score, breakpoint localization using boundary bias, and overall copy number accuracy using root mean squared error (RMSE). A predicted CNA was considered a true positive if it was of the same type as a ground-truth CNA and overlapped at least 20% of the ground-truth event length. Unmatched predicted events were considered false positives, and unmatched ground-truth events were considered false negatives. Precision, recall, and F1 score were calculated as

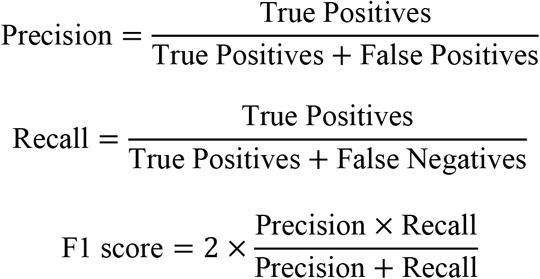

For each true-positive event with ground-truth interval [*e*_1_, *e*_2_] and predicted interval [*ê*_1_, *ê*_2_], boundary bias (Mb) was defined as |*ê*_1_ − *e*_1_| +|*ê*_2_ − *e*_2_|, which is the sum of the absolute distances between the predicted and true breakpoints at the two boundaries. Boundary bias was averaged across true-positive events within each cell, and cells with no true-positive events were excluded from this metric. Overall copy number accuracy was quantified for each cell using the bin-level RMSE between predicted and ground-truth states,

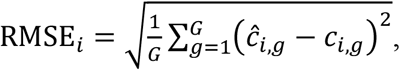

where *ĉ*_*i,g*_ and *c*_*i,g*_ denote the predicted and ground-truth states, respectively, for cell *i* at genomic bin *g*.

### Application to a colorectal cancer single-cell multi-omics dataset

We analyzed patient CRC01 from a human colorectal cancer cohort profiled using scTrio-seq2 (Bian et al. 2018), which simultaneously measures the genome and transcriptome of the same single cells. CRC01 was microsatellite-stable and classified as the consensus molecular subtype CMS2, which is characterized by frequent CNAs (Bian et al. 2018; Guinney et al. 2015). Because some cells were profiled for only one modality, we retained cells with both DNA and RNA measurements after quality control. All sequencing data were aligned to the GRCh37/hg19 reference genome.

Because the genomic component of scTrio-seq2 is based on single-cell bisulfite sequencing, we used total sequencing coverage within each bin for CNA inference. Raw sequencing reads were trimmed using Trim Galore (Krueger et al. 2021) to remove adapters and low-quality bases, aligned using Bismark (Krueger and Andrews 2011), and deduplicated using SAMtools (Li et al. 2009). Deduplicated reads were counted across 25,262 non-overlapping 100 kb genomic bins and normalized using normal cells as the diploid reference. For the transcriptome, reads were trimmed using cutadapt (Martin 2011), aligned using STAR (Dobin et al. 2013), and quantified per gene using RSEM (Li and Dewey 2011). Genes detected in fewer than 5% of cells were removed, retaining 13,855 of 54,849 genes. Gene-expression signals were then processed as described in Data preprocessing and mapped to the same 100 kb genomic bins as the DNA data, resulting in 11,632 bins with RNA measurements.

For the matched bulk WGS dataset, paired-end reads were aligned using BWA (Li and Durbin 2009), sorted and marked for duplicates, and merged across sequencing lanes of each tumor site with Picard (Broad Institute 2019). Read depth was calculated over the same 100 kb genomic bins using mosdepth (Pedersen and Quinlan 2018), normalized against the matched blood sample, and segmented into copy number states using DNAcopy (Olshen et al. 2004). Because single-cell ground-truth CNA profiles were unavailable, the matched bulk WGS profile from each tumor site was used as an orthogonal reference. Concordance between each inferred single-cell CNA profile and the bulk WGS profile was quantified using Pearson correlation and RMSE.

### Application to an IDH-mutant glioma 10x Multiome dataset

We analyzed single-nucleus 10x Multiome data from the CARE IDH-mutant glioma cohort (Johnson et al. 2026), a longitudinal study of clonal and cell state evolution across matched primary and recurrent resections. We restricted the analysis to samples with at least 1,900 ATAC nuclei passing quality control and matched bulk WGS, resulting in 20 samples (14 oligodendrogliomas and 6 astrocytomas) from 10 patients (Supplementary Fig. 7). All data were aligned to the GRCh38/hg38 reference genome.

Cell type annotations were obtained from the original study, which annotated nuclei in both the RNA and ATAC modalities (Johnson et al. 2026). Myeloid, endothelial and lymphocyte nuclei were used to construct the diploid reference. Oligodendrocyte and astrocyte nuclei were excluded from the reference to minimize potential contamination by misannotated malignant nuclei carrying the 1p/19q codeletion. ATAC fragments were counted in 500 kb bins, and bins with mappability < 0.50, GC content outside the range 0.20–0.80, or zero fragments in > 85% of nuclei were excluded. *L2R* values were calculated using the pooled diploid reference profile. RNA data were restricted to nuclei retained after ATAC quality control and processed as described in Data preprocessing.

Reference copy number profiles were obtained from matched bulk WGS. Segmented WGS calls were lifted from GRCh37 to GRCh38 coordinates with UCSC liftOver (Hinrichs et al. 2006) and mapped to the same 500 kb genomic bins. We benchmarked MintCNA against four scATAC-seq CNA callers: epiAneufinder (Ramakrishnan et al. 2023), AtaCNA (Wang et al. 2025), Copy-scAT (Nikolic et al. 2021) and RIDDLER (Moore et al. 2026). The benchmark also included the scRNA-seq caller inferCNV (Patel et al. 2014), and CONGAS+ (Patruno et al. 2023) which jointly models scATAC-seq and scRNA-seq. For comparison with bulk WGS, the output of each method was converted to a pseudobulk *L2R* profile by averaging across malignant nuclei within each 500 kb bin (Supplementary Methods). Performance was evaluated using the Pearson correlation between each inferred pseudobulk profile and the matched bulk WGS profile, together with bin-level precision, recall and F1 score.

Agreement between subclone labels and RNA expression was assessed by leave-one-out 15-nearest-neighbor classification in RNA principal component space. Differential expression between MintCNA-defined subclones was tested for genes expressed in at least 5% of the 995 malignant nuclei in sample P111T2 using a two-sided Wilcoxon rank-sum test, with Benjamini-Hochberg correction for multiple testing. Promoter accessibility was quantified within 2 kb of each transcription start site, and differences in promoter accessibility between subclones were compared with corresponding expression changes using Spearman correlation. Motif accessibility within the same regions was quantified using chromVAR (Schep et al. 2017) for 633 JASPAR2020 motifs (Fornes et al. 2020) and compared between subclones. Among the genes and motifs evaluated, 22 marker genes associated with OPC-like and AC-like states (Neftel et al. 2019) and motifs of 12 lineage transcription factors (Rowitch and Kriegstein 2010) were highlighted to characterize lineage identity.

For both real data applications, all methods were run with default settings unless otherwise specified (Supplementary Methods) and all analyses were restricted to autosomes.

## Data availability

The colorectal cancer scTrio-seq2 data are available from the NCBI Gene Expression Omnibus (GEO) under accession GSE97693 (https://www.ncbi.nlm.nih.gov/geo/query/acc.cgi?acc=GSE97693). The matched WGS data are available under controlled access from the European Genome-phenome Archive (EGA) under accession EGAS00001003242 (https://ega-archive.org/studies/EGAS00001003242). Access to these data can be requested through the EGA data access committee. The IDH-mutant glioma 10x Multiome datasets from the CARE cohort are available from the GEO under accessions GSE327580 (ATAC, https://www.ncbi.nlm.nih.gov/geo/query/acc.cgi?acc=GSE327580) and GSE326221 (RNA, https://www.ncbi.nlm.nih.gov/geo/query/acc.cgi?acc=GSE326221). The matched bulk WGS data and cell-type annotations are available through Synapse under project syn63600501 (https://www.synapse.org/care_idh_mutant). The simulated scDNA-seq and paired scRNA-seq data generated in this study have been deposited in Zenodo (https://doi.org/10.5281/zenodo.23385650).

## Code availability

MintCNA is available as open source and available on GitHub (https://github.com/WENHANB/MintCNA) under the GPL-3.0 license.

## Acknowledgements

This work was supported in part by the University of Florida Artificial Intelligence Initiative (to F.X.).

## Author contributions

W.B. developed the method, implemented the software, designed and performed the simulations and benchmarks, analyzed the colorectal cancer and glioma datasets, and wrote the original draft of the manuscript. F.X. conceived and supervised the study, guided the methodological development, and revised the manuscript in detail. All authors read and approved the final manuscript.

## References

Baslan T, Hicks J. 2017. Unravelling biology and shifting paradigms in cancer with single-cell sequencing. Nat Rev Cancer 17: 557–569.

Beroukhim R, Mermel CH, Porter D, Wei G, Raychaudhuri S, Donovan J, Barretina J, Boehm JS, Dobson J, Urashima M, et al. 2010. The landscape of somatic copy-number alteration across human cancers. Nature 463: 899–905.

Bhattacharya A, Bense RD, Urzúa-Traslaviña CG, De Vries EGE, Van Vugt MATM, Fehrmann RSN. 2020. Transcriptional effects of copy number alterations in a large set of human cancers. Nat Commun 11: 715.

Bian S, Hou Y, Zhou X, Li X, Yong J, Wang Y, Wang W, Yan J, Hu B, Guo H, et al. 2018. Single-cell multiomics sequencing and analyses of human colorectal cancer. Science 362: 1060–1063.

Broad Institute. 2019. Picard toolkit. http://broadinstitute.github.io/picard/.

Chen S, Lake BB, Zhang K. 2019. High-throughput sequencing of the transcriptome and chromatin accessibility in the same cell. Nat Biotechnol 37: 1452–1457.

De Falco A, Caruso F, Su X-D, Iavarone A, Ceccarelli M. 2023. A variational algorithm to detect the clonal copy number substructure of tumors from scRNA-seq data. Nat Commun 14: 1074.

Dempster AP, Laird NM, Rubin DB. 1977. Maximum Likelihood from Incomplete Data Via the EM Algorithm. Royal Statistical Society Journal Series B: Methodological 39: 1–22.

Dobin A, Davis CA, Schlesinger F, Drenkow J, Zaleski C, Jha S, Batut P, Chaisson M, Gingeras TR. 2013. STAR: ultrafast universal RNA-seq aligner. Bioinformatics 29: 15–21.

Dong X, Zhang L, Hao X, Wang T, Vijg J. 2020. SCCNV: A Software Tool for Identifying Copy Number Variation From Single-Cell Whole-Genome Sequencing. Front Genet 11: 505441.

Follain B, Wang T, Samworth RJ. 2022. High-dimensional Changepoint Estimation with Heterogeneous Missingness. Journal of the Royal Statistical Society Series B: Statistical Methodology 84: 1023–1055.

Fornes O, Castro-Mondragon JA, Khan A, van der Lee R, Zhang X, Richmond PA, Modi BP, Correard S, Gheorghe M, Baranašić D, et al. 2020. JASPAR 2020: update of the open-access database of transcription factor binding profiles. Nucleic Acids Res 48: D87–D92.

Gao R, Bai S, Henderson YC, Lin Y, Schalck A, Yan Y, Kumar T, Hu M, Sei E, Davis A, et al. 2021. Delineating copy number and clonal substructure in human tumors from single-cell transcriptomes. Nat Biotechnol 39: 599–608.

Garvin T, Aboukhalil R, Kendall J, Baslan T, Atwal GS, Hicks J, Wigler M, Schatz MC. 2015. Interactive analysis and assessment of single-cell copy-number variations. Nat Methods 12: 1058–1060.

Glasgow SM, Zhu W, Stolt CC, Huang T-W, Chen F, LoTurco JJ, Neul JL, Wegner M, Mohila C, Deneen B. 2014. Mutual antagonism between Sox10 and NFIA regulates diversification of glial lineages and glioma subtypes. Nat Neurosci 17: 1322–1329.

Guinney J, Dienstmann R, Wang X, de Reyniès A, Schlicker A, Soneson C, Marisa L, Roepman P, Nyamundanda G, Angelino P, et al. 2015. The consensus molecular subtypes of colorectal cancer. Nat Med 21: 1350–1356.

Hinrichs AS, Karolchik D, Baertsch R, Barber GP, Bejerano G, Clawson H, Diekhans M, Furey TS, Harte RA, Hsu F, et al. 2006. The UCSC Genome Browser Database: update 2006. Nucleic Acids Res 34: D590–D598.

Hui S, Nielsen R. 2022. SCONCE: a method for profiling copy number alterations in cancer evolution using single-cell whole genome sequencing ed. C. Alkan. Bioinformatics 38: 1801–1808.

Jamal-Hanjani M, Quezada SA, Larkin J, Swanton C. 2015. Translational Implications of Tumor Heterogeneity. Clin Cancer Res 21: 1258–1266.

Johnson KC, Spitzer A, Varn FS, Nomura M, Garofano L, Chowdhury T, Lipsa A, Zhang L, Fernández EC, Barak T, et al. 2026. Acquired genetic and cell-state changes in IDH-mutant glioma progression. Nature 655: 1048–1059.

Krueger F, Andrews SR. 2011. Bismark: a flexible aligner and methylation caller for Bisulfite-Seq applications. Bioinformatics 27: 1571–1572.

Krueger F, James F, Ewels P, Afyounian E, Schuster-Boeckler B. 2021. FelixKrueger/TrimGalore: v0.6.7 - DOI via Zenodo. https://zenodo.org/records/5127899 (Accessed April 10, 2026).

Kuipers J, Tuncel MA, Ferreira PF, Jahn K, Beerenwinkel N. 2025. Single-cell copy number calling and event history reconstruction ed. R. Schwartz. Bioinformatics 41: btaf072.

Li B, Dewey CN. 2011. RSEM: accurate transcript quantification from RNA-Seq data with or without a reference genome. BMC Bioinformatics 12: 323.

Li H, Durbin R. 2009. Fast and accurate short read alignment with Burrows–Wheeler transform. Bioinformatics 25: 1754–1760.

Li H, Handsaker B, Wysoker A, Fennell T, Ruan J, Homer N, Marth G, Abecasis G, Durbin R, 1000 Genome Project Data Processing Subgroup. 2009. The Sequence Alignment/Map format and SAMtools. Bioinformatics 25: 2078–2079.

Li R, Alberge J-B, Keshavarzian T, Tsuji J, Gustafsson J, Rahmat M, Lightbody ED, Deng SL, Riviero S, Miller M, et al. 2025a. Numbat-multiome: inferring copy number variations by combining RNA and chromatin accessibility information from single-cell data. Brief Bioinform 26: bbaf516.

Li S, Alexander J, Kendall J, Andrews P, Rose E, Orjuela H, Park S, Podszus C, Shanley L, Ranade N, et al. 2025b. Hybrid BAG-seq: DNA and RNA from the same single nucleus reveals interactions between genomic and transcriptomic landscapes in human tumor samples. Genome Biol 26: 314.

Liu F, Shi F, Yu Z. 2024a. Inferring single-cell copy number profiles through cross-cell segmentation of read counts. BMC Genomics 25: 25.

Liu R, Hu H, Zhang S, Deng Y, Liu Z, Chen Y, Chen Z. 2024b. An ECG denoising technique based on AHIN block and gradient difference max loss. Journal of Electrocardiology 86: 153761.

Louis DN, Perry A, Wesseling P, Brat DJ, Cree IA, Figarella-Branger D, Hawkins C, Ng HK, Pfister SM, Reifenberger G, et al. 2021. The 2021 WHO Classification of Tumors of the Central Nervous System: a summary. Neuro Oncol 23: 1231–1251.

Ma S, Zhang B, LaFave LM, Earl AS, Chiang Z, Hu Y, Ding J, Brack A, Kartha VK, Tay T, et al. 2020. Chromatin Potential Identified by Shared Single-Cell Profiling of RNA and Chromatin. Cell 183: 1103-1116.e20.

Magi A, Tattini L, Cifola I, D’Aurizio R, Benelli M, Mangano E, Battaglia C, Bonora E, Kurg A, Seri M, et al. 2013. EXCAVATOR: detecting copy number variants from whole-exome sequencing data. Genome Biol 14: R120.

Mallory XF, Edrisi M, Navin N, Nakhleh L. 2020. Methods for copy number aberration detection from single-cell DNA-sequencing data. Genome Biol 21: 208.

Mamlouk S, Childs LH, Aust D, Heim D, Melching F, Oliveira C, Wolf T, Durek P, Schumacher D, Bläker H, et al. 2017. DNA copy number changes define spatial patterns of heterogeneity in colorectal cancer. Nat Commun 8: 14093.

Martin M. 2011. Cutadapt removes adapter sequences from high-throughput sequencing reads. EMBnet.journal 17: 10–12.

Mathieu M, Couprie C, LeCun Y. 2016. Deep multi-scale video prediction beyond mean square error. http://arxiv.org/abs/1511.05440 (Accessed June 17, 2026).

Moore TW, Mohammed H, Adey AC, Yardimci GG. 2026. Robust and generalizable CNV detection for single-cell sequencing assays. Nucleic Acids Res 54: gkag636.

Navin N, Kendall J, Troge J, Andrews P, Rodgers L, McIndoo J, Cook K, Stepansky A, Levy D, Esposito D, et al. 2011. Tumour evolution inferred by single-cell sequencing. Nature 472: 90–94.

Neftel C, Laffy J, Filbin MG, Hara T, Shore ME, Rahme GJ, Richman AR, Silverbush D, Shaw ML, Hebert CM, et al. 2019. An Integrative Model of Cellular States, Plasticity, and Genetics for Glioblastoma. Cell 178: 835-849.e21.

Nikolic A, Singhal D, Ellestad K, Johnston M, Shen Y, Gillmor A, Morrissy S, Cairncross JG, Jones S, Lupien M, et al. 2021. Copy-scAT: Deconvoluting single-cell chromatin accessibility of genetic subclones in cancer. SCIENCE ADVANCES.

Olsen TR, Talla P, Sagatelian RK, Furnari J, Bruce JN, Canoll P, Zha S, Sims PA. 2025. Scalable co-sequencing of RNA and DNA from individual nuclei. Nat Methods 22: 477–487.

Olshen AB, Venkatraman ES, Lucito R, Wigler M. 2004. Circular binary segmentation for the analysis of array-based DNA copy number data. Biostatistics 5: 557–572.

Patel AP, Tirosh I, Trombetta JJ, Shalek AK, Gillespie SM, Wakimoto H, Cahill DP, Nahed BV, Curry WT, Martuza RL, et al. 2014. Single-cell RNA-seq highlights intratumoral heterogeneity in primary glioblastoma. Science 344: 1396–1401.

Patruno L, Milite S, Bergamin R, Calonaci N, D’Onofrio A, Anselmi F, Antoniotti M, Graudenzi A, Caravagna G. 2023. A Bayesian method to infer copy number clones from single-cell RNA and ATAC sequencing ed. T.M. Przytycka. PLoS Comput Biol 19: e1011557.

Pedersen BS, Quinlan AR. 2018. Mosdepth: quick coverage calculation for genomes and exomes. Bioinformatics 34: 867–868.

Pino MS, Chung DC. 2010. The Chromosomal Instability Pathway in Colon Cancer. Gastroenterology 138: 2059–2072.

Ramakrishnan A, Symeonidi A, Hanel P, Schmid KT, Richter ML, Schubert M, Colomé-Tatché M. 2023. epiAneufinder identifies copy number alterations from single-cell ATAC-seq data. Nat Commun 14: 5846.

Reyal F, Stransky N, Bernard-Pierrot I, Vincent-Salomon A, de Rycke Y, Elvin P, Cassidy A, Graham A, Spraggon C, Désille Y, et al. 2005. Visualizing Chromosomes as Transcriptome Correlation Maps: Evidence of Chromosomal Domains Containing Co-expressed Genes— A Study of 130 Invasive Ductal Breast Carcinomas. Cancer Res 65: 1376–1383.

Rowitch DH, Kriegstein AR. 2010. Developmental genetics of vertebrate glial–cell specification. Nature 468: 214–222.

Salari K, Spulak ME, Cuff J, Forster AD, Giacomini CP, Huang S, Ko ME, Lin AY, van de Rijn M, Pollack JR. 2012. CDX2 is an amplified lineage-survival oncogene in colorectal cancer. Proceedings of the National Academy of Sciences 109: E3196–E3205.

Schep AN, Wu B, Buenrostro JD, Greenleaf WJ. 2017. chromVAR: inferring transcription-factor-associated accessibility from single-cell epigenomic data. Nat Methods 14: 975–978.

Shah SP, Xuan X, DeLeeuw RJ, Khojasteh M, Lam WL, Ng R, Murphy KP. 2006. Integrating copy number polymorphisms into array CGH analysis using a robust HMM. Bioinformatics 22: e431–e439.

Sheffer M, Bacolod MD, Zuk O, Giardina SF, Pincas H, Barany F, Paty PB, Gerald WL, Notterman DA, Domany E. 2009. Association of survival and disease progression with chromosomal instability: A genomic exploration of colorectal cancer. Proceedings of the National Academy of Sciences 106: 7131–7136.

Siegel RL, Wagle NS, Cercek A, Smith RA, Jemal A. 2023. Colorectal cancer statistics, 2023. CA: A Cancer Journal for Clinicians 73: 233–254.

Stratton MR. 2011. Exploring the Genomes of Cancer Cells: Progress and Promise. Science 331: 1553–1558.

The Cancer Genome Atlas Network. 2012. Comprehensive molecular characterization of human colon and rectal cancer. Nature 487: 330–337.

Vogelstein B, Papadopoulos N, Velculescu VE, Zhou S, Diaz LA, Kinzler KW. 2013. Cancer Genome Landscapes. Science 339: 1546–1558.

Wang R, Lin D-Y, Jiang Y. 2020. SCOPE: A Normalization and Copy-Number Estimation Method for Single-Cell DNA Sequencing. Cell Systems 10: 445-452.e6.

Wang R, Zhang Y, Wang M, Feng X, Wang J, Li SC. 2022. Resolving single-cell copy number profiling for large datasets. Briefings in Bioinformatics 23: bbac264.

Wang X, Jin Z, Shi Y, Xi R. 2025. Detecting copy-number alterations from single-cell chromatin sequencing data by AtaCNA. Cell Reports Methods 5: 100939.

Weiner S, Bansal MS. 2023. CNAsim: improved simulation of single-cell copy number profiles and DNA-seq data from tumors ed. C. Kendziorski. Bioinformatics 39: btad434.

Woo S, Park J, Lee J-Y, Kweon IS. 2018. CBAM: Convolutional Block Attention Module. In Computer Vision – ECCV 2018 (eds. V. Ferrari, M. Hebert, C. Sminchisescu, and Y. Weiss), Vol. 11211 of Lecture Notes in Computer Science, pp. 3–19, Springer International Publishing, Cham https://link.springer.com/10.1007/978-3-030-01234-2_1 (Accessed December 11, 2025).

Yu L, Wang X, Mu Q, Tam SST, Loi DSC, Chan AKY, Poon WS, Ng H-K, Chan DTM, Wang J, et al. 2023a. scONE-seq: A single-cell multi-omics method enables simultaneous dissection of phenotype and genotype heterogeneity from frozen tumors. Sci Adv 9: eabp8901.

Yu Z, Liu F, Shi F, Du F. 2023b. rcCAE: a convolutional autoencoder method for detecting intratumor heterogeneity and single-cell copy number alterations. Briefings in Bioinformatics 24: bbad108.

Zack TI, Schumacher SE, Carter SL, Cherniack AD, Saksena G, Tabak B, Lawrence MS, Zhang C-Z, Wala J, Mermel CH, et al. 2013. Pan-cancer patterns of somatic copy number alteration. Nat Genet 45: 1134–1140.

Zappia L, Phipson B, Oshlack A. 2017. Splatter: simulation of single-cell RNA sequencing data. Genome Biol 18: 174.

Zhao Y, Luquette LJ, Veit AD, Wang X, Xi R, Viswanadham VV, Zhang Y, Shao DD, Walsh CA, Yang HW, et al. 2025. High-resolution detection of copy number alterations in single cells with HiScanner. Nat Commun 16: 5477.

