## Supplementary Information for "MintCNA: A Unified Framework for Integrative Copy Number Profiling with Single-Cell Multi-Omics Data"

Wenhan Bao[1], Feifei Xiao[1*]

[1] Department of Biostatistics, College of Public Health and Health Professions and College of Medicine, University of Florida, Gainesville, FL, 32603, USA

[*] To whom correspondence should be addressed

Corresponding to:

Feifei Xiao, Ph.D.

Department of Biostatistics, College of Public Health and Health Professions and College of Medicine, University of Florida

2004 Mowry Rd., CTRB building, Room 5227, Gainesville, FL, 32603

### 1. Supplementary Methods

#### Attention-guided denoising convolutional autoencoder (AG-DCAE)

*Network architecture and activation functions*

AG-DCAE is a one-dimensional convolutional autoencoder for denoising the coverage-based *L2R* profiles (i.e., $x_i$). The encoder $f_\theta(\cdot)$ consists of three stacked 1D convolutional layers with channel sizes $1 \rightarrow 128 \rightarrow 64 \rightarrow 32$, a kernel size of 5 and a stride of 1, each followed by LeakyReLU activation. A convolutional block attention module (CBAM; Woo et al. 2018) with reduction ratio 4 is applied to the feature map of the final convolutional layer to adaptively emphasize informative channels and genomic positions. Let $F_i$ denote this feature map for cell $i$ and $F_i^{att}$ the output of CBAM. The two are combined through a gated residual connection $\widetilde{F_i} = F_i + \beta(F_i^{att} - F_i)$ with $\beta = 0.1$. The attention-refined feature map $\widetilde{F_i}$ is flattened and passed through a fully connected bottleneck with LeakyReLU activation to obtain the latent representation $Z_i \in \mathbb{R}^q$ with $q = 250$. The decoder $d_\theta(\cdot)$ is symmetric to the encoder, with a fully connected layer followed by three 1D transposed-convolutional layers with channel sizes $32 \rightarrow 64 \rightarrow 128 \rightarrow 1$ and LeakyReLU activations between layers. All convolutional and fully connected weights are initialized from $\mathcal{N}(0, 0.02^2)$ with zero biases.

*Training procedure*

During training, additive Gaussian noise $\eta_i \sim \mathcal{N}(0, \sigma_\eta^2 I)$ with $\sigma_\eta^2 = 0.08$ is added to the input profile $x_i$ and the network is trained to reconstruct $x_i$ from the noisy input by minimizing $\mathcal{L}(\theta)$ (Methods) with $\lambda_1 = 0.2$. Parameters are optimized using Adam (Kingma and Ba 2017) with learning rate $1 \times 10^{-4}$ and a mini-batch size of 16. Cells are split into training (80%) and validation (20%) sets. Early stopping with a patience of 15 epochs and a minimum improvement of 0.001 is applied to the validation loss, and the model with the lowest validation loss is used as the final model.

*Subclone identification*

Subclones are identified by $k$-means clustering (Lloyd 1982) of the latent representations $Z_i$ of all cells. Prior to clustering, latent representations are standardized and projected onto the top principal components explaining at least 95% of the variance (capped at 20 components) to reduce noise in the latent space. The number of clusters is selected by a stability-based approach (Monti et al. 2003). For each candidate number of clusters, k-means is applied to 200 random 80% subsamples of cells, and the adjusted Rand index (ARI) is computed between pairs of subsampled solutions. The number of clusters with the highest mean pairwise ARI is selected, and a one-standard-error rule (Hastie et al. 2009) is applied to choose fewer clusters when multiple values have similar stability. The final clustering is obtained by running k-means on all cells with 200 random initializations.

**Segmentation and change-point detection**

*Quantile normalization*

To alleviate potential violations of the normality assumption, we apply a quantile-normal transformation to each omics layer independently across the genome. For each cell $i$ and layer $k$, observed denoised *L2R* values $\widehat{X_{\iota,k,g}}$ are ranked and mapped to a normal distribution by $\tilde{X}_{i,k,g} = \hat{\sigma}_k \cdot \Phi^{-1}((r_{i,k,g} - 0.5)/n_{i,k})$, where $r_{i,k,g}$ is the rank of bin $g$ among the $n_{i,k}$ observed bins of layer $k$ (i.e., $\Omega_{i,k,g} = 1$) and $\Phi^{-1}$ is the standard normal quantile function. We set $\hat{\sigma}_k$ to the median standard deviation of layer $k$ across all cells, making all cells share a common scale for each layer.

*Closed-form alternating solution*

The $\ell_2$-regularized rank-1 approximation is solved by alternating updates. For a given window centered at $g$, let $\hat{v}^{(t)}$ and $\hat{w}^{(t)}$ denote the estimates of $v$ and $w$ at iteration $t$. The procedure is initialized with $\hat{v}^{(0)}$ set to the leading left singular vector of $T^{(g)}$. Given the current omics-weight vector $\hat{v}^{(t)}$, the within-window weight vector is updated as

$$\hat{w}^{(t+1)} = \frac{T^{(g)\top}\hat{v}^{(t)}}{\| T^{(g)\top}\hat{v}^{(t)} \|_2},$$

which is the projection of $T^{(g)\top}\hat{v}^{(t)}$ onto the unit sphere. Given the current $\hat{w}^{(t+1)}$, the omics-weight vector is updated as

$$\hat{v}^{(t+1)} = \frac{1}{2\lambda_2} T^{(g)}\hat{w}^{(t+1)},$$

which is the closed-form ridge solution. The procedure alternates between these two steps until the change in $\hat{v}^{(t)}$ falls below $10^{-10}$ or 1,000 iterations are reached. At convergence, the estimates are taken as $\hat{v}^{(g)}$ and $\hat{w}^{(g)}$ (Methods).

*Choice of the tuning parameter $\lambda_2$*

Each row of $T^{(g)}$ is the local CUSUM statistics of one omics layer, computed over the observed bins only. We set the ridge parameter to

$$\lambda_2 = \frac{1}{2}\sqrt{P \log(K \log P)},$$

adapted from Follain et al. (2022), where $P$ is the number of bins in the chromosome being scanned and $K$ is the number of omics layers.

*Significance thresholding*

To assess the significance of candidate breakpoints while controlling for multiple comparisons (Hao et al. 2013), we estimate the null distribution of $| D(g) |$ under the assumption of no change point. Null data are generated as simulated replicates with a fixed random seed. For each omics layer, bin values are drawn independently from $N(0, \hat{\sigma}_k^2)$, and the missingness pattern $\Omega_i$ is preserved. Each replicate is scanned chromosome by chromosome. Local maxima of $|D(g)|$ within $\pm 2h$ bins are collected, and when two peaks lie closer than $h$ bins, the earlier one is discarded. Peak heights are pooled over chromosomes and replicates until 10,000 peaks have been collected or all replicates have been scanned, and they form the empirical null distribution. Candidate positions (i.e., the local maxima of $|D(g)|$ within $\pm h$ bins) are identified as breakpoints when $|D(g)| > \tau_\alpha$, where $\tau_\alpha$ is the $(1-\alpha)$ quantile of this null distribution. By default, we use $h = 5$ bins, $\alpha = 0.05$ and 1 null replicate.

**CNA state assignment**

*Cross-omics segment fusion*

For each cell $i$, the detected breakpoints $\hat{g}_{i,1}^* < \cdots < \hat{g}_{i,M_i-1}^*$ partition the genome into $M_i$ contiguous segments. Let segment $s$ be the index set

$$J_{i,s} = \{\hat{g}_{i,s-1}^* + 1, \dots, \hat{g}_{i,s}^*\}, s = 1, \dots, M_i,$$

where $\hat{g}_{i,0}^* = 0$ and $\hat{g}_{i,M_i}^* = G$. Given the quantile-normalized *L2R* matrix $\tilde{X}_i$ and observation indicator $\Omega_i$, we compute omics-specific segment means

$$\bar{x}_{i,k,s} = \frac{1}{n_{i,k,s}} \sum_{g \in J_{i,s}} \Omega_{i,k,g} \tilde{X}_{i,k,g}, n_{i,k,s} = \sum_{g \in J_{i,s}} \Omega_{i,k,g},$$

where $n_{i,k,s}$ is the number of observed bins in segment $s$ for omics layer $k$. We then combine the segment means across omics layers to obtain a fused segment mean

$$Y_{i,s} = \sum_{k: n_{i,k,s} > 0} u_{i,k,s} \bar{x}_{i,k,s},$$

where the weights are proportional to the numbers of observed bins in the segment

$$u_{i,k,s} = \frac{n_{i,k,s}}{\sum_{k': n_{i,k',s} > 0} n_{i,k',s}}.$$

In the single-omics setting ($K = 1$), $Y_{i,s}$ reduces to the segment mean of the coverage-based layer.

*Expectation–maximization algorithm for truncated Gaussian mixture model*

MintCNA assigns each segment $s$ of cell $i$ to one of $C$ CNA states, with $C = 3$ (deletion, neutral and duplication) or $C = 5$ (deletion, loss, neutral, gain and amplification). Let $z_{i,s} \in \{1, \ldots, C\}$ denote the latent CNA state for segment $s$. We model the fused segment mean $Y_{i,s}$ by a truncated Gaussian mixture

$$Y_{i,s} \mid z_{i,s} = c \sim TN(\mu_c, \sigma_c^2; a_c, b_c), \Pr(z_{i,s} = c) = \pi_c,$$

where $TN(\mu_c, \sigma_c^2; a_c, b_c)$ denotes the normal distribution $N(\mu_c, \sigma_c^2)$ truncated to $[a_c, b_c]$, $\pi_c > 0$ and $\sum_{c=1}^{C} \pi_c = 1$. The truncation bounds $[a_c, b_c]$ keep the states in order and are fixed on the quantile-normalized scale. For $C = 5$, they are

$$(-\infty, -\delta_2],\ (-\delta_2, -\delta_1],\ (-\delta_1, \delta_1],\ (\delta_1, \delta_2],\ (\delta_2, \infty),$$

and for $C = 3$, they are

$$(-\infty, -\delta_1],\ (-\delta_1, \delta_1],\ (\delta_1, \infty),$$

where $0 < \delta_1 < \delta_2$ are fixed constants, with $\delta_1 = 0.30$ and $\delta_2 = 0.5$ by default.

Let $\phi(\cdot)$ and $\Phi(\cdot)$ denote the probability density function and cumulative distribution function of the standard normal distribution. The truncated Gaussian density of state $c$ is

$$f_c(y; \mu_c, \sigma_c^2, a_c, b_c) = \frac{1}{\sigma_c} \frac{\phi\left(\frac{y - \mu_c}{\sigma_c}\right)}{\Phi\left(\frac{b_c - \mu_c}{\sigma_c}\right) - \Phi\left(\frac{a_c - \mu_c}{\sigma_c}\right)} 1\{a_c \le y \le b_c\}.$$

For each cell $i$, the observed-data log-likelihood is

$$l(\Theta) = \sum_{s=1}^{M_i} \log\left(\sum_{c=1}^{C} \pi_c\, f_c\big(Y_{i,s}; \mu_c, \sigma_c^2, a_c, b_c\big)\right), \Theta = \{(\pi_c, \mu_c, \sigma_c^2)\}_{c=1}^{C}.$$

**EM updates.** Starting from empirically initialized parameters $\hat{\Theta}^{(0)}$, the EM algorithm iterates between the E-step and the M-step and produces a sequence of estimates $\hat{\Theta}^{(t)}$, $t = 0,1,2,\ldots$.

**E-step.** Given $\hat{\Theta}^{(t)}$, we compute the posterior probability that segment $s$ of cell $i$ is in state $c$

$$\gamma_{i,s,c}^{(t+1)} = \frac{\hat{\pi}_c^{(t)} f_c\left(Y_{i,s}; \hat{\mu}_c^{(t)}, \left(\hat{\sigma}_c^{(t)}\right)^2, a_c, b_c\right)}{\sum_{c'=1}^{C} \hat{\pi}_{c'}^{(t)} f_{c'}\left(Y_{i,s}; \hat{\mu}_{c'}^{(t)}, \left(\hat{\sigma}_{c'}^{(t)}\right)^2, a_{c'}, b_{c'}\right)}.$$

**M-step.** We update $(\pi_c, \mu_c, \sigma_c^2)$ using weighted moments

$$n_c^{(t+1)} = \sum_{s=1}^{M_i} \gamma_{i,s,c}^{(t+1)}, \quad \hat{\pi}_c^{(t+1)} = \frac{n_c^{(t+1)}}{M_i}, \quad \hat{\mu}_c^{(t+1)} = \frac{1}{n_c^{(t+1)}} \sum_{s=1}^{M_i} \gamma_{i,s,c}^{(t+1)} Y_{i,s},$$

$$\left(\hat{\sigma}_c^{(t+1)}\right)^2 = \max\left\{\frac{1}{n_c^{(t+1)}} \sum_{s=1}^{M_i} \gamma_{i,s,c}^{(t+1)} \left(Y_{i,s} - \hat{\mu}_c^{(t+1)}\right)^2,\ \sigma_{\min}^2\right\},$$

where $n_c^{(t+1)}$ is the expected number of segments in state $c$, and $\sigma_{\min}^2 = 10^{-3}$ is a lower bound that keeps the variances from shrinking to zero. The algorithm stops after 30 iterations or earlier when the summed change in the state means falls below $10^{-6}$.

The final CNA state of segment $s$ in cell $i$ is assigned by maximum posterior probability

$$\hat{z}_{i,s} = \arg\max_{c \in \{1,\ldots,C\}} \gamma_{i,s,c},$$

where $\gamma_{i,s,c}$ is the posterior probability at convergence. Adjacent segments with the same assigned state are merged. In the implementation, the variance of each $Y_{i,s}$ is scaled by $1/|J_{i,s}|$ and the moments in the M-step are weighted by $|J_{i,s}|$, so that longer segments receive more weight.

**Single-cell sequencing data simulation**

Ground-truth single-cell CNA profiles were generated using CNAsim (Weiner and Bansal 2023), which simulates a cell-lineage tree with divergent subclonal populations and evolves CNAs along its branches. We used the direct read count simulation mode (mode 1), in which the read count

$\mathrm{RC}_{i,g}$ of cell $i$ at bin $g$ is drawn from $\mathrm{Poisson}(\kappa \cdot \mathrm{CN}_{i,g} \cdot \omega_g)$ where $\mathrm{CN}_{i,g}$ is the underlying copy number, $\kappa$ is the scale factor corresponding to sequencing coverage and $\omega_g$ models uneven amplification across bins. This uneven amplification $\omega_g$ was modeled by a beta distribution defined by the point (0.5, 0.4) on the Lorenz curve. To simulate technical variability, each bin-level read count was multiplied by Gaussian noise $\xi_{i,g} \sim N(1, \nu^2)$ with $\nu \in \{0, 0.05, 0.1\}$. All simulations used the 22 autosomes of the hg38 reference genome with 100 kb bins.

The cell population and its CNAs were simulated with the same parameters at all coverages and noise levels. Each simulated dataset contained 100 cells, including 10 normal cells without CNAs, and three tumor subclones. The number of focal CNAs on each branch of the cell-lineage tree followed a Poisson distribution with mean 6. The branch leading to the founder tumor cell carried 10 times as many focal CNAs, which became clonal CNAs shared by all tumor cells. Focal CNAs had a mean length of 5 Mb and a minimum length of 10 kb. Chromosome-level CNAs had a mean of two events on the branch leading to the founder tumor cell and one on each branch leading to a subclone. Each chromosome-level CNA affected one chromosome arm (probability 0.75) or the whole chromosome (probability 0.25). For both focal and chromosome-level CNAs, gains and losses were equally likely, and no whole-genome duplication was simulated.

**Other CNA detection benchmarking methods**

For the simulation and colorectal cancer benchmarks, we compared MintCNA with six scDNA-seq CNA callers. All methods were run according to their documentation, with default settings when available, and the scripts are available in the MintCNA repository.

HMMcopy version 1.46.0 (Shah et al. 2006) is an R-based read-depth caller that corrects for GC content and mappability and segments with a hidden Markov model. We applied `correctReadcount` and then `HMMsegment` with the long-segment parameters from the HMMcopy vignette (e = 0.9999999 and strength = 1e7).

SCOPE version 1.16.0 (Wang et al. 2020) is an R-based method that uses a Poisson latent factor model for normalization and cross-sample segmentation. We normalized read counts with `normalize_scope_foreach` using the known normal cells, kept the fit with the largest Bayesian information criterion and segmented each chromosome with `segment_CBScs`.

SCCNV (Dong et al. 2020) corrects sequencing biases and calls CNAs from read depth across genomic bins. We re-implemented its published workflow in R, which filters low-quality bins, normalizes read counts, smooths them with a sliding window and computes posterior probabilities for copy number states 0–20.

SCONCE (Hui and Nielsen 2022) is a hidden Markov model-based caller that uses normal cells as negative controls. We built the diploid reference from the normal cells with the package scripts and ran `sconce` on each tumor cell with a maximum ploidy of 5.

rcCAE (Yu et al. 2023) combines a convolutional autoencoder with latent-space clustering and hidden Markov model-based CNA calling. We trained the autoencoder with default settings and called CNAs with `SCHMM` in MATLAB with a maximum copy number of 6.

SCICoNE version 1.0.0 (Kuipers et al. 2025) reconstructs CNA profiles by jointly modeling CNA events and clonal evolution in a phylogenetic framework. We ran `breakpoint_detection` and then `inference` with default settings.

For the glioma multiome benchmark, all methods were applied to the quality-filtered fragment files and gene expression matrices for the 20 samples from the source study, restricted to the annotated barcodes. Methods requiring normal reference cells used the same diploid reference nuclei as MintCNA (Methods). Each method was run using its published default settings. Only the modifications required to run each method on these data are described below.

epiAneufinder version 1.1.5 (Ramakrishnan et al. 2023) bins scATAC-seq fragment counts, corrects them for GC content and segments each cell into losses, gains and normal regions. We ran it on each sample with the documented settings (windowSize = 1e5, minFrags = 20000, and k = 4) and excluded the sex chromosomes and the mitochondrial genome.

AtaCNA version 0.1.0 (Wang et al. 2025) normalizes each cell against non-tumor cells and jointly segments all cells with the multi-sample BIC-seq algorithm. We ran it in normal-cell mode with default settings. Because `generate_input()` did not complete on multiome-scale files, we assigned fragments to the package's 1-Mb bins by their midpoints, which yields the same count matrix.

Copy-scAT version 0.40 (Nikolic et al. 2021) calls arm-level gains and losses by fitting Gaussian mixture models to CpG-normalized fragment coverage summarized over chromosome arms. We followed the authors' tutorial with the published parameter values and the main workflow, without the experimental normal-cell option.

RIDDLER (Moore et al. 2026) models the expected coverage of each genomic window from GC content, mappability and accessibility peaks and calls CNAs from significant deviations. We ran it at its published 1-Mb resolution with at least 10,000 fragments per cell.

inferCNV version 1.26.0 (Patel et al. 2014) detects CNAs in scRNA-seq data by comparing smoothed expression along each chromosome between malignant and reference cells. We ran it on the raw counts of malignant and reference nuclei with gene coordinates from EnsDb.Hsapiens.v86 and default settings (cutoff = 1 and denoise = FALSE).

CONGAS+ version 0.2.0 (Patruno et al. 2023) is a Bayesian model of scRNA-seq and scATAC-seq counts over genomic segments that identifies copy number clones. We ran it in multiome mode with negative binomial likelihoods and equal modality weights (lambdas = 0.5). CONGAS+ requires a sample-specific segmentation, but the only one available was the bulk WGS reference used for evaluation. We therefore used the package's arm-level segmentation (`hg38_arms`) without the acrocentric short arms 13p, 14p, 15p, 21p, and 22p, which contain almost no reads.

**Evaluation of the benchmark results with the glioma multiome dataset**

To evaluate each method against bulk WGS, its output was averaged over malignant nuclei in each 500 kb bin and converted to pseudobulk *L2R* relative to its neutral value. Outputs given per chromosome arm or per gene were first assigned to the overlapping bins, and outputs given per clone were assigned to the nuclei of that clone. Copy-scAT arms that were tested but not called were treated as neutral. The Pearson correlation was calculated to evaluate the concordance between the pseudobulk and bulk WGS profiles of each sample (Fig. 5a). Following AtaCNA (Wang et al. 2025), each nucleus in each bin was labeled as loss (i.e., copy ratio < 0.75), gain (i.e., > 1.25) or neutral. For precision and recall (Fig. 5b), each bin was labeled loss, neutral or gain by the most common state among malignant nuclei. A bin was a true positive when this label had the same direction as the bulk WGS call, and counts were pooled across the 20 samples.

**References**


Dong X, Zhang L, Hao X, Wang T, Vijg J. 2020. SCCNV: A Software Tool for Identifying Copy Number Variation From Single-Cell Whole-Genome Sequencing. *Front Genet* **11**: 505441.

Follain B, Wang T, Samworth RJ. 2022. High-dimensional Changepoint Estimation with Heterogeneous Missingness. *Journal of the Royal Statistical Society Series B: Statistical Methodology* **84**: 1023–1055.

Hao N, Selena Niu Y, Zhang H. 2013. Multiple change-point detection via a screening and ranking algorithm. *STAT SINICA*. http://www3.stat.sinica.edu.tw/statistica/J23N4/J23N47/J23N47.html (Accessed December 9, 2025).

Hastie T, Tibshirani R, Friedman J. 2009. *The Elements of Statistical Learning*. Springer New York, New York, NY http://link.springer.com/10.1007/978-0-387-84858-7 (Accessed March 17, 2026).

Hui S, Nielsen R. 2022. SCONCE: a method for profiling copy number alterations in cancer evolution using single-cell whole genome sequencing ed. C. Alkan. *Bioinformatics* **38**: 1801–1808.

Kingma DP, Ba J. 2017. Adam: A Method for Stochastic Optimization. http://arxiv.org/abs/1412.6980 (Accessed December 11, 2025).

Kuipers J, Tuncel MA, Ferreira PF, Jahn K, Beerenwinkel N. 2025. Single-cell copy number calling and event history reconstruction ed. R. Schwartz. *Bioinformatics* **41**: btaf072.

Lloyd S. 1982. Least squares quantization in PCM. *IEEE Trans Inform Theory* **28**: 129–137.

Monti S, Tamayo P, Mesirov J, Golub T. 2003. Consensus Clustering: A Resampling-Based Method for Class Discovery and Visualization of Gene Expression Microarray Data. *Machine Learning* **52**: 91–118.

Moore TW, Mohammed H, Adey AC, Yardımcı GG. 2026. Robust and generalizable CNV detection for single-cell sequencing assays. *Nucleic Acids Res* **54**: gkag636.

Nikolic A, Singhal D, Ellestad K, Johnston M, Shen Y, Gillmor A, Morrissy S, Cairncross JG, Jones S, Lupien M, et al. 2021. Copy-scAT: Deconvoluting single-cell chromatin accessibility of genetic subclones in cancer. *SCIENCE ADVANCES*.

Patel AP, Tirosh I, Trombetta JJ, Shalek AK, Gillespie SM, Wakimoto H, Cahill DP, Nahed BV, Curry WT, Martuza RL, et al. 2014. Single-cell RNA-seq highlights intratumoral heterogeneity in primary glioblastoma. *Science* **344**: 1396–1401.

Patruno L, Milite S, Bergamin R, Calonaci N, D'Onofrio A, Anselmi F, Antoniotti M, Graudenzi A, Caravagna G. 2023. A Bayesian method to infer copy number clones from single-cell RNA and ATAC sequencing ed. T.M. Przytycka. *PLoS Comput Biol* **19**: e1011557.

Ramakrishnan A, Symeonidi A, Hanel P, Schmid KT, Richter ML, Schubert M, Colomé-Tatché M. 2023. epiAneufinder identifies copy number alterations from single-cell ATAC-seq data. *Nat Commun* **14**: 5846.

Shah SP, Xuan X, DeLeeuw RJ, Khojasteh M, Lam WL, Ng R, Murphy KP. 2006. Integrating copy number polymorphisms into array CGH analysis using a robust HMM. *Bioinformatics* **22**: e431–e439.

Wang R, Lin D-Y, Jiang Y. 2020. SCOPE: A Normalization and Copy-Number Estimation Method for Single-Cell DNA Sequencing. *Cell Systems* **10**: 445-452.e6.

Wang X, Jin Z, Shi Y, Xi R. 2025. Detecting copy-number alterations from single-cell chromatin sequencing data by AtaCNA. *Cell Reports Methods* **5**: 100939.

Weiner S, Bansal MS. 2023. CNAsim: improved simulation of single-cell copy number profiles and DNA-seq data from tumors ed. C. Kendziorski. *Bioinformatics* **39**: btad434.

Woo S, Park J, Lee J-Y, Kweon IS. 2018. CBAM: Convolutional Block Attention Module. http://arxiv.org/abs/1807.06521 (Accessed December 11, 2025).

Yu Z, Liu F, Shi F, Du F. 2023. rcCAE: a convolutional autoencoder method for detecting intra-tumor heterogeneity and single-cell copy number alterations. *Briefings in Bioinformatics* **24**: bbad108.

#### 2. Supplementary Figures

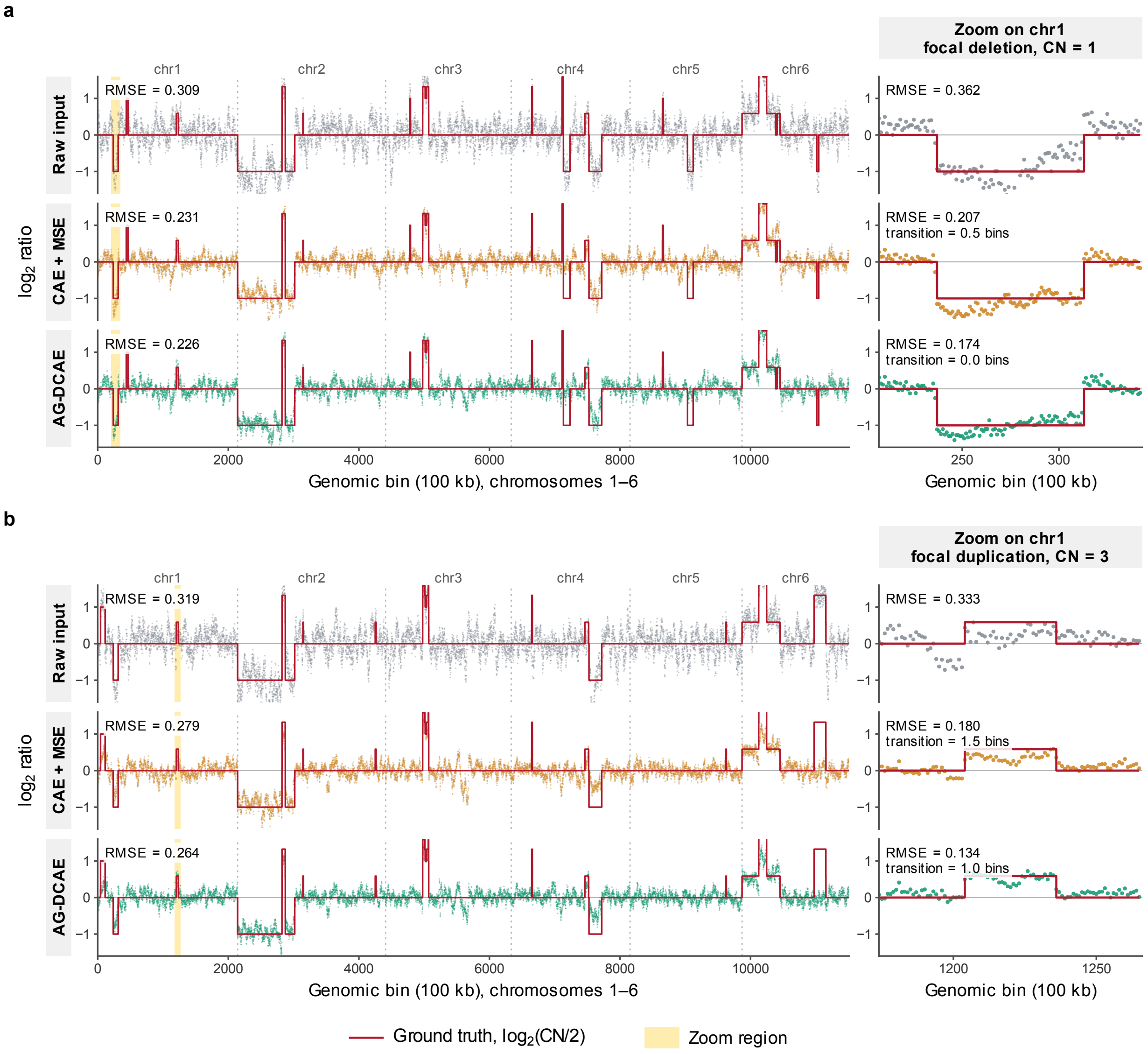


**Supplementary Figure 1. AG-DCAE reduces noise while preserving CNA boundaries.** Two simulated cells generated by CNAsim at 0.1× coverage with a noise level of 0.1. Rows show the raw *L2R* profile (top), the CAE + MSE baseline without attention or edge-preserving loss (middle) and AG-DCAE with edge-preserving loss (bottom). Left column displays chromosomes 1–6 with the zoom region shaded in yellow. Right column shows the focal CNA. Red step function indicates ground-truth log2(CN/2). (a) Focal deletion (CN = 1) on chromosome 1. (b) Focal duplication (CN = 3) on chromosome 1. RMSE: root mean squared error between each profile and the ground truth across the bins shown. Transition, mean number of bins within 8 bins of a true breakpoint whose denoised value lies in the central 50% band between the two flanking denoised levels (lower values indicate sharper boundaries).

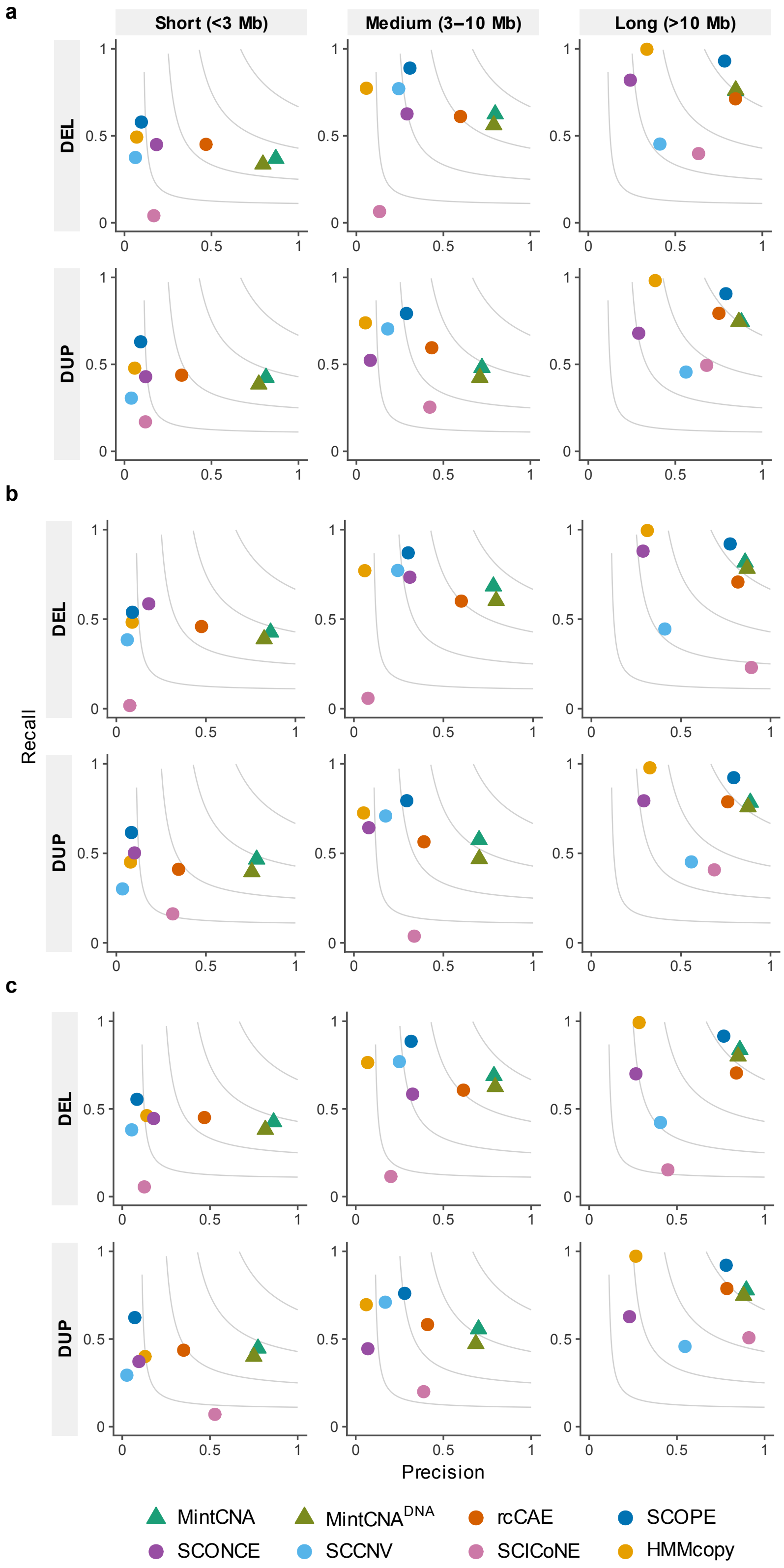


**Supplementary Figure 2. Event-level detection accuracy stratified by CNA size at 0.1× coverage.** Precision-recall plots for deletions (upper rows) and duplications (lower rows) across three event size categories (<3 Mb, 3–10 Mb, >10 Mb). Panels a-c correspond to noise levels of 0, 0.05, and 0.1, respectively. F1 contour lines at 0.2, 0.4, 0.6 and 0.8 are shown in gray.

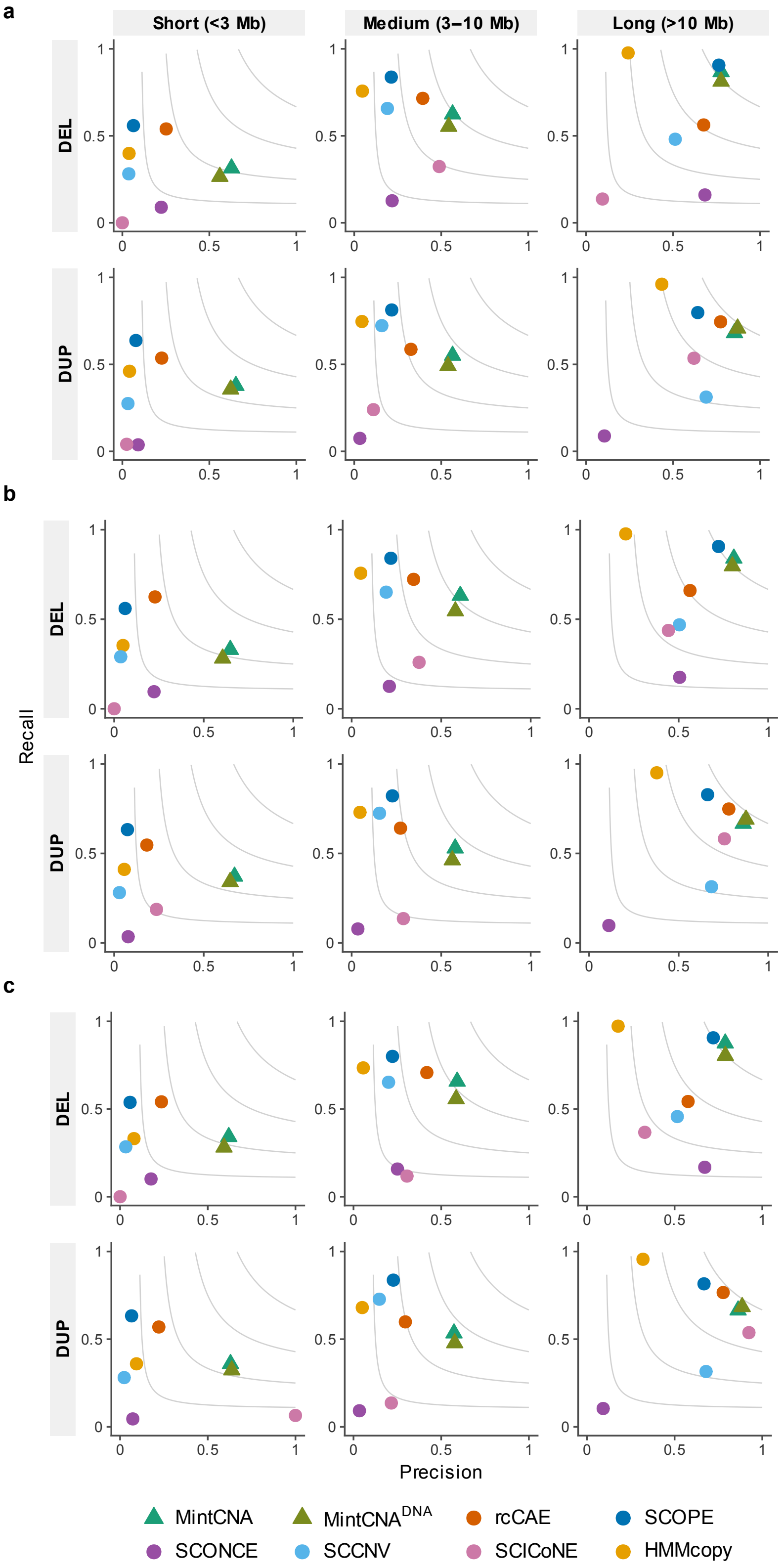


**Supplementary Figure 3. Event-level detection accuracy stratified by CNA size at 0.05× coverage.** Precision-recall plots for deletions (upper rows) and duplications (lower rows) across three event size categories (<3 Mb, 3–10 Mb, >10 Mb). Panels a-c correspond to noise levels of 0, 0.05, and 0.1, respectively. F1 contour lines at 0.2, 0.4, 0.6 and 0.8 are shown in gray.

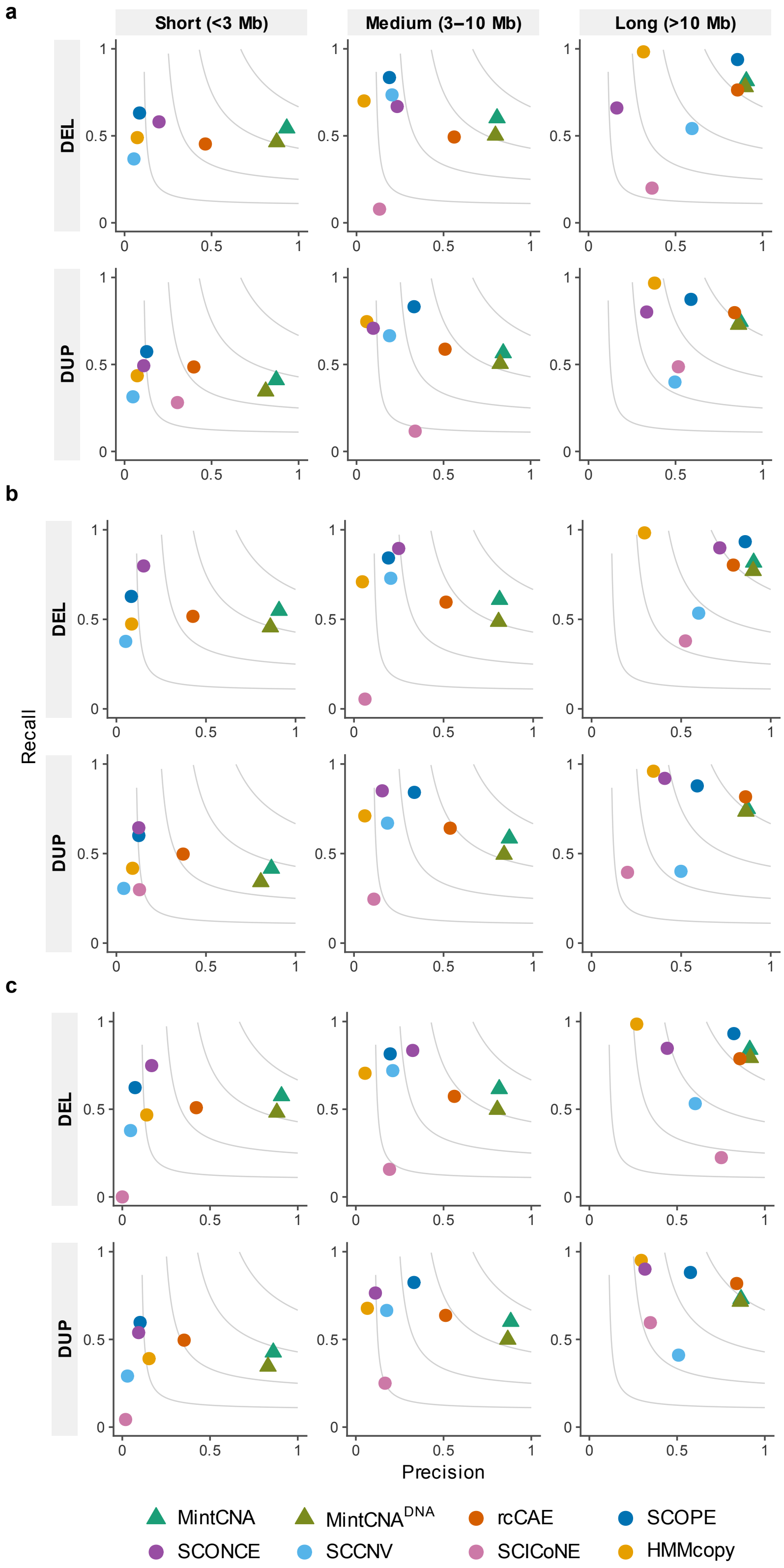


**Supplementary Figure 4. Event-level detection accuracy stratified by CNA size at 0.2× coverage.** Precision-recall plots for deletions (upper rows) and duplications (lower rows) across three event size categories (<3 Mb, 3–10 Mb, >10 Mb). Panels a-c correspond to noise levels of 0, 0.05, and 0.1, respectively. F1 contour lines at 0.2, 0.4, 0.6 and 0.8 are shown in gray.

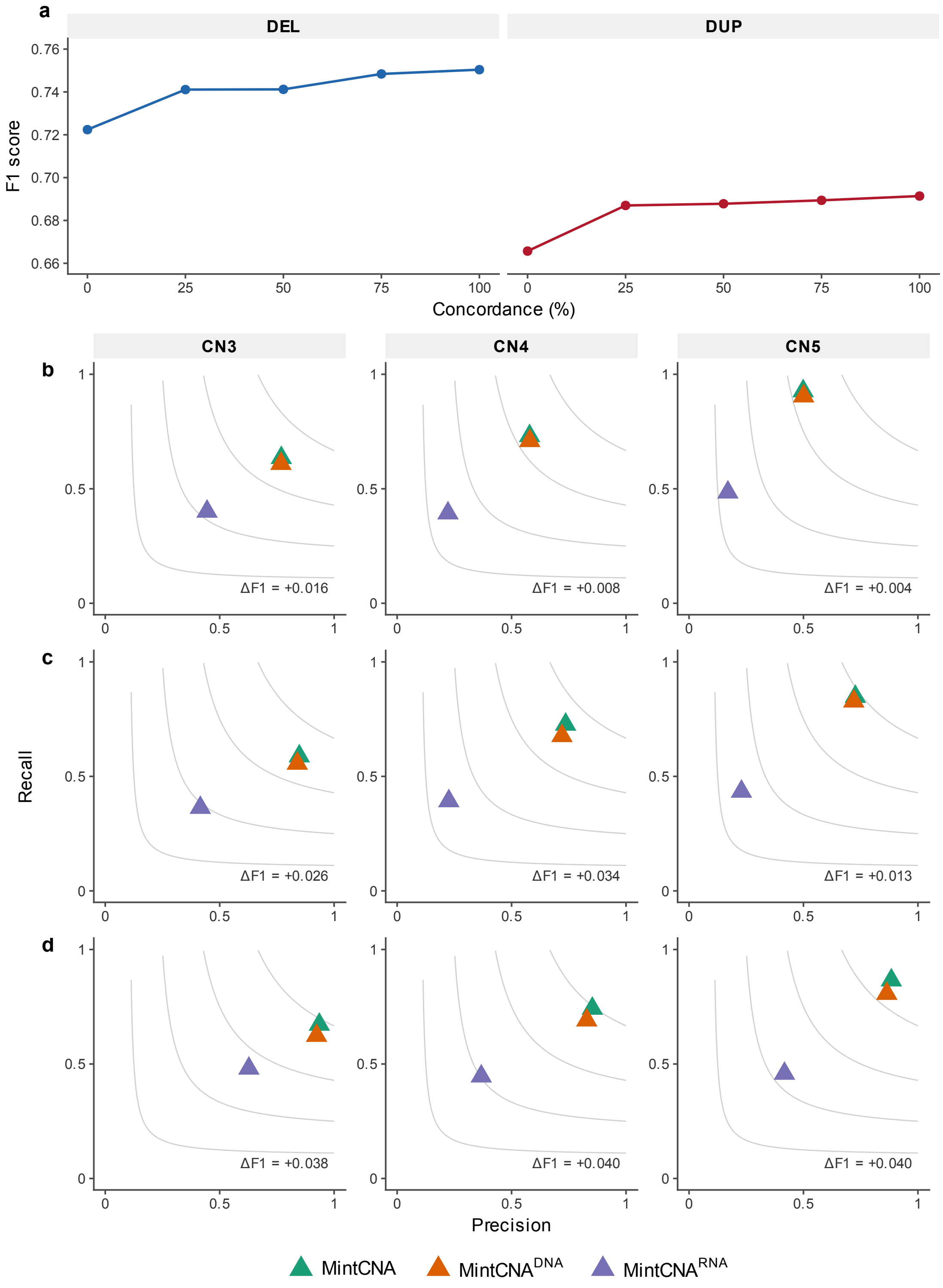


**Supplementary Figure 5. Multi-omics integration improves CNA detection across concordance levels and copy numbers.** (a) F1 score of MintCNA at varying concordance levels for deletions (left) and duplications (right) under the baseline simulation (0.1× coverage, no noise). Concordance: the proportion of genes whose expression reflects the underlying copy number state. (b–d) Precision-recall comparison among MintCNA, MintCNA$^{DNA}$ using scDNA-seq data alone, and MintCNA$^{RNA}$ using scRNA-seq data alone, stratified by copy number (CN=3, CN=4, CN≥5) at coverage 0.05× (b), 0.1× (c), and 0.2× (d) with no additional noise. ΔF1: the F1 score difference between MintCNA and MintCNA$^{DNA}$.

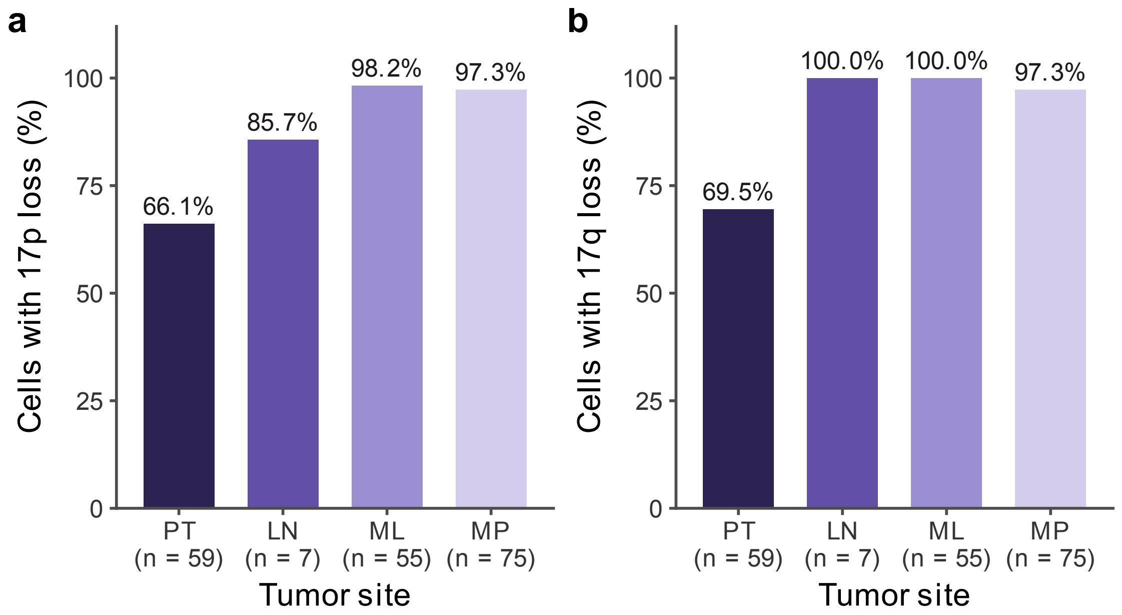


**Supplementary Figure 6. Proportion of cells with chromosome 17 arm-level loss across tumor sites of patient CRC01.** Percentage of cells classified as harboring loss on chromosome 17p (a) and 17q (b) in the primary tumor (PT), lymph node metastasis (LN), liver metastasis (ML), and post-treatment liver metastasis (MP). A bin-level majority rule was applied, where a chromosome arm was classified as lost in a cell if > 50% of its bins were called as loss or deletion by MintCNA. $n$: number of cells at each site.

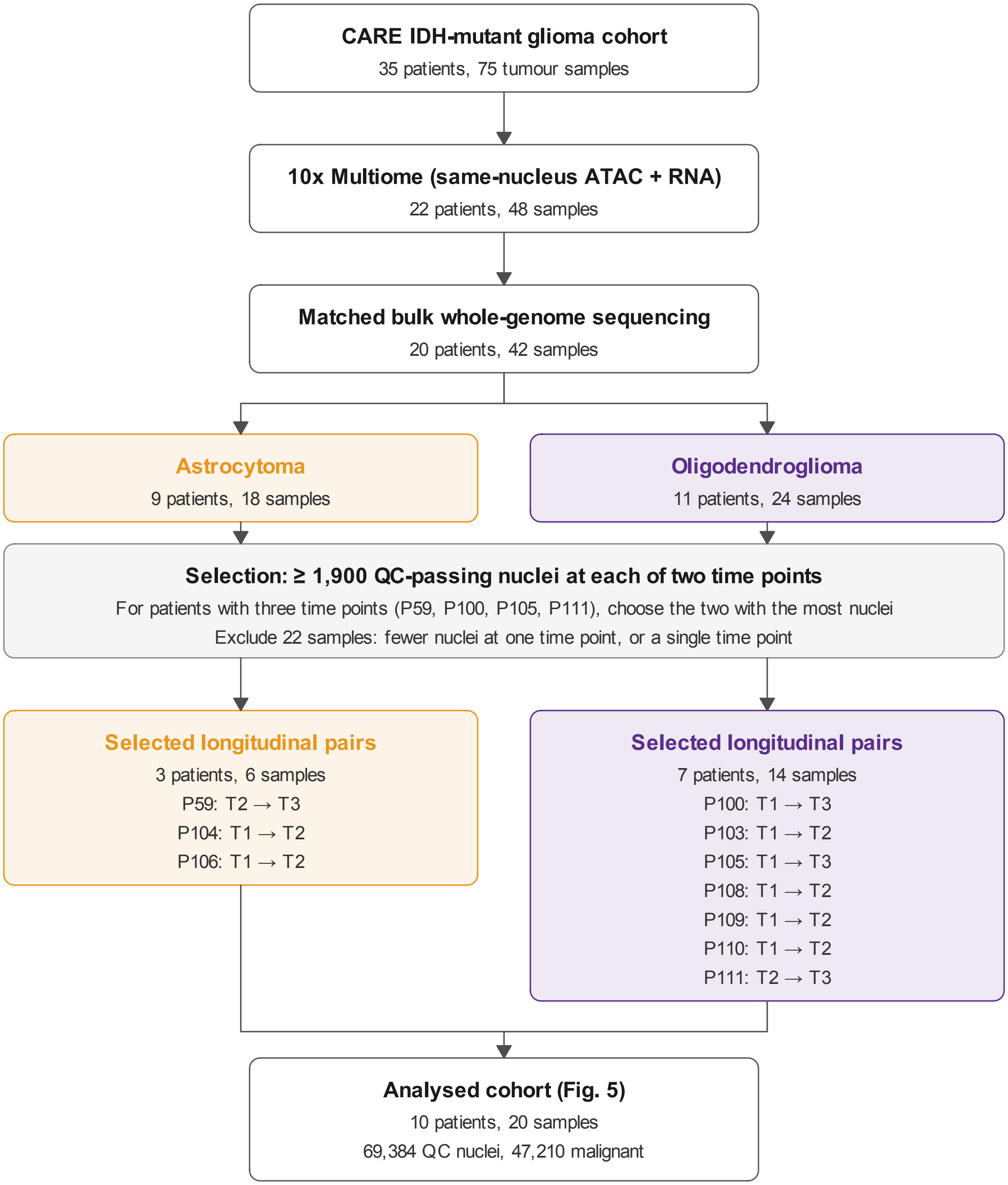


**Supplementary Figure 7. Sample selection from the CARE glioma cohort.** Of 75 tumor samples (35 patients) in the CARE IDH-mutant glioma cohort, 48 were profiled by 10x Multiome among which 42 had matched bulk whole-genome sequencing. Patients were retained when two time points each with > 1,900 nuclei passing quality control. For the patients with three time points, the two time points with the most nuclei were used. T1, T2 and T3 denote the initial tumor, first recurrence and second recurrence.

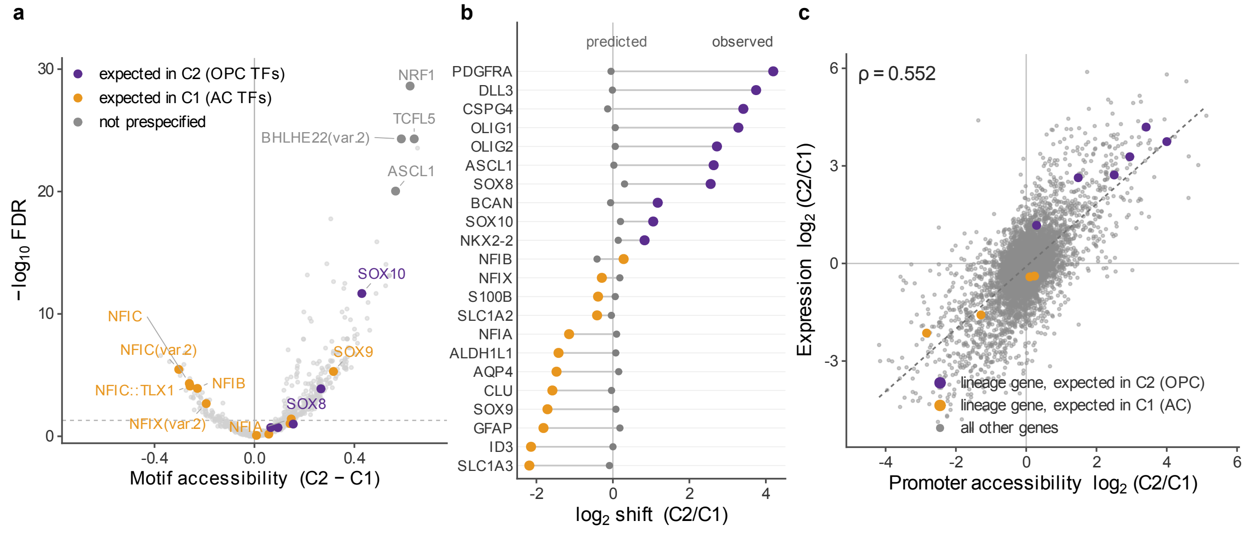


**Supplementary Figure 8. Motif accessibility and gene dosage in the subclones of sample P111T2.** (a) Difference in accessibility of 633 JASPAR2020 motifs between subclones C2 (OPC-like) and C1 (AC-like). Motifs of 12 prespecified lineage transcription factors are colored by the subclone in which they are expected to be more accessible. The FDR threshold was set at 0.05. (b) Expression changes of 22 lineage genes, predicted from the copy number difference between subclones (gray) and observed (colored). (c) Correlation between promoter accessibility change and gene expression change between subclones for 6,317 genes with the same copy number in both subclones (i.e., absolute difference in mean copy number < 0.1).

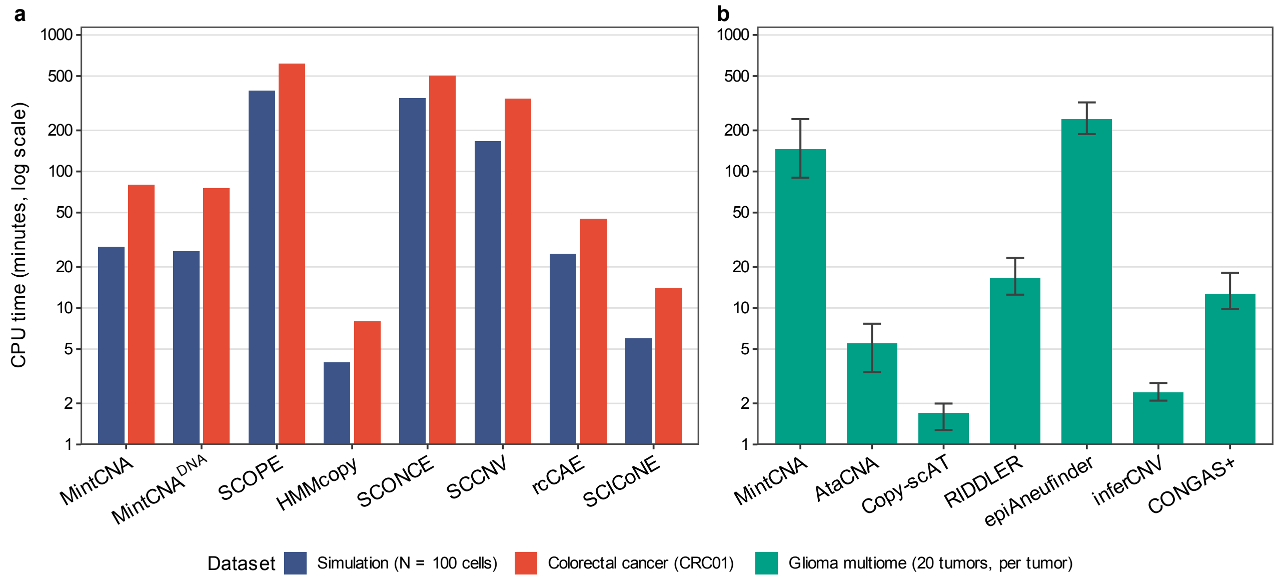


**Supplementary Figure 9. Computational time of CNA detection methods.** (a) Paired scDNA-seq and scRNA-seq benchmark. Each method was run on the simulated dataset (N = 100 cells) and on the colorectal cancer scTrio-seq2 dataset (CRC01) using 8 CPU cores and 64 GB RAM. (b) Benchmark on the glioma 10x Multiome dataset. CPU time per tumor was calculated as total core-minutes to enable comparison across methods using different numbers of CPU cores. Bars indicate medians and whiskers show interquartile ranges across 20 samples.

#### 3. Supplementary Tables

**Supplementary Table 1. Event-level detection accuracy for deletions across simulation scenarios.** Precision, recall and F1 score of each method under three coverage depths (0.05×, 0.1× and 0.2×) and three noise levels (0, 0.05 and 0.1), with 90 tumor cells per scenario. A predicted deletion (CN < 2) was counted as a true positive if it overlapped at least 20% of a ground-truth deletion. MintCNA$^{DNA}$: the DNA-only mode of MintCNA.

| Coverage | Noise | Method | Precision | Recall | F1 |
|---|---|---|---|---|---|
| 0.05 | 0 | MintCNA | 0.692 | 0.588 | 0.636 |
| 0.05 | 0 | MintCNA$^{DNA}$ | 0.665 | 0.526 | 0.588 |
| 0.05 | 0 | rcCAE | 0.395 | 0.710 | 0.507 |
| 0.05 | 0 | HMMcopy | 0.079 | 0.926 | 0.145 |
| 0.05 | 0 | SCOPE | 0.155 | 0.858 | 0.262 |
| 0.05 | 0 | SCONCE | 0.286 | 0.129 | 0.178 |
| 0.05 | 0 | SCCNV | 0.132 | 0.634 | 0.218 |
| 0.05 | 0 | SCICoNE | 0.307 | 0.170 | 0.219 |
| 0.05 | 0.05 | MintCNA | 0.714 | 0.582 | 0.642 |
| 0.05 | 0.05 | MintCNA$^{DNA}$ | 0.695 | 0.520 | 0.595 |
| 0.05 | 0.05 | rcCAE | 0.354 | 0.784 | 0.487 |
| 0.05 | 0.05 | HMMcopy | 0.091 | 0.915 | 0.166 |
| 0.05 | 0.05 | SCOPE | 0.150 | 0.853 | 0.255 |
| 0.05 | 0.05 | SCONCE | 0.286 | 0.139 | 0.187 |
| 0.05 | 0.05 | SCCNV | 0.129 | 0.634 | 0.214 |
| 0.05 | 0.05 | SCICoNE | 0.331 | 0.186 | 0.238 |
| 0.05 | 0.1 | MintCNA | 0.693 | 0.606 | 0.647 |
| 0.05 | 0.1 | MintCNA$^{DNA}$ | 0.689 | 0.524 | 0.595 |
| 0.05 | 0.1 | rcCAE | 0.400 | 0.732 | 0.518 |
| 0.05 | 0.1 | HMMcopy | 0.110 | 0.897 | 0.196 |
| 0.05 | 0.1 | SCOPE | 0.146 | 0.833 | 0.248 |
| 0.05 | 0.1 | SCONCE | 0.279 | 0.152 | 0.197 |
| 0.05 | 0.1 | SCCNV | 0.121 | 0.631 | 0.202 |
| 0.05 | 0.1 | SCICoNE | 0.356 | 0.159 | 0.220 |
| 0.1 | 0 | MintCNA | 0.918 | 0.614 | 0.736 |
| 0.1 | 0 | MintCNA$^{DNA}$ | 0.903 | 0.578 | 0.705 |
| 0.1 | 0 | rcCAE | 0.653 | 0.628 | 0.641 |
| 0.1 | 0 | HMMcopy | 0.108 | 0.959 | 0.195 |
| 0.1 | 0 | SCOPE | 0.237 | 0.883 | 0.374 |
| 0.1 | 0 | SCONCE | 0.304 | 0.768 | 0.436 |
| 0.1 | 0 | SCCNV | 0.189 | 0.723 | 0.300 |
| 0.1 | 0 | SCICoNE | 0.303 | 0.127 | 0.179 |
| 0.1 | 0.05 | MintCNA | 0.889 | 0.664 | 0.760 |
| 0.1 | 0.05 | MintCNA$^{DNA}$ | 0.891 | 0.604 | 0.720 |
| 0.1 | 0.05 | rcCAE | 0.655 | 0.628 | 0.642 |
| 0.1 | 0.05 | HMMcopy | 0.119 | 0.951 | 0.212 |
| 0.1 | 0.05 | SCOPE | 0.234 | 0.872 | 0.369 |
| 0.1 | 0.05 | SCONCE | 0.303 | 0.860 | 0.449 |
| 0.1 | 0.05 | SCCNV | 0.184 | 0.724 | 0.293 |
| 0.1 | 0.05 | SCICoNE | 0.253 | 0.113 | 0.156 |
| 0.1 | 0.1 | MintCNA | 0.893 | 0.669 | 0.765 |
| 0.1 | 0.1 | MintCNA$^{DNA}$ | 0.891 | 0.621 | 0.732 |
| 0.1 | 0.1 | rcCAE | 0.658 | 0.623 | 0.640 |
| 0.1 | 0.1 | HMMcopy | 0.142 | 0.936 | 0.247 |
| 0.1 | 0.1 | SCOPE | 0.223 | 0.879 | 0.356 |

| Coverage | Noise | Method | Precision | Recall | F1 |
|---|---|---|---|---|---|
| 0.1 | 0.1 | SCONCE | 0.305 | 0.682 | 0.421 |
| 0.1 | 0.1 | SCCNV | 0.171 | 0.716 | 0.276 |
| 0.1 | 0.1 | SCICoNE | 0.290 | 0.137 | 0.187 |
| 0.2 | 0 | MintCNA | 0.911 | 0.658 | 0.764 |
| 0.2 | 0 | $\text{MintCNA}^{\text{DNA}}$ | 0.900 | 0.585 | 0.709 |
| 0.2 | 0 | rcCAE | 0.633 | 0.581 | 0.606 |
| 0.2 | 0 | HMMcopy | 0.099 | 0.946 | 0.179 |
| 0.2 | 0 | SCOPE | 0.188 | 0.887 | 0.311 |
| 0.2 | 0 | SCONCE | 0.235 | 0.746 | 0.357 |
| 0.2 | 0 | SCCNV | 0.173 | 0.721 | 0.280 |
| 0.2 | 0 | SCICoNE | 0.224 | 0.081 | 0.119 |
| 0.2 | 0.05 | MintCNA | 0.904 | 0.663 | 0.765 |
| 0.2 | 0.05 | $\text{MintCNA}^{\text{DNA}}$ | 0.899 | 0.573 | 0.700 |
| 0.2 | 0.05 | rcCAE | 0.585 | 0.669 | 0.624 |
| 0.2 | 0.05 | HMMcopy | 0.108 | 0.943 | 0.193 |
| 0.2 | 0.05 | SCOPE | 0.186 | 0.889 | 0.308 |
| 0.2 | 0.05 | SCONCE | 0.260 | 0.949 | 0.408 |
| 0.2 | 0.05 | SCCNV | 0.169 | 0.723 | 0.273 |
| 0.2 | 0.05 | SCICoNE | 0.251 | 0.130 | 0.172 |
| 0.2 | 0.1 | MintCNA | 0.907 | 0.679 | 0.777 |
| 0.2 | 0.1 | $\text{MintCNA}^{\text{DNA}}$ | 0.903 | 0.587 | 0.712 |
| 0.2 | 0.1 | rcCAE | 0.613 | 0.653 | 0.633 |
| 0.2 | 0.1 | HMMcopy | 0.127 | 0.936 | 0.224 |
| 0.2 | 0.1 | SCOPE | 0.174 | 0.883 | 0.291 |
| 0.2 | 0.1 | SCONCE | 0.294 | 0.918 | 0.445 |
| 0.2 | 0.1 | SCCNV | 0.159 | 0.720 | 0.260 |
| 0.2 | 0.1 | SCICoNE | 0.313 | 0.124 | 0.178 |

**Supplementary Table 2. Event-level detection accuracy for duplications across simulation scenarios.** Precision, recall and F1 score of each method under three coverage depths (0.05×, 0.1× and 0.2×) and three noise levels (0, 0.05 and 0.1), with 90 tumor cells per scenario. A predicted duplication (CN > 2) was counted as a true positive if it overlapped at least 20% of a ground-truth duplication. $MintCNA^{DNA}$: the DNA-only mode of MintCNA.

| Coverage | Noise | Method | Precision | Recall | F1 |
|---|---|---|---|---|---|
| 0.05 | 0 | MintCNA | 0.731 | 0.567 | 0.639 |
| 0.05 | 0 | $MintCNA^{DNA}$ | 0.729 | 0.552 | 0.628 |
| 0.05 | 0 | rcCAE | 0.397 | 0.695 | 0.506 |
| 0.05 | 0 | HMMcopy | 0.090 | 0.917 | 0.164 |
| 0.05 | 0 | SCOPE | 0.187 | 0.861 | 0.307 |
| 0.05 | 0 | SCONCE | 0.095 | 0.110 | 0.102 |
| 0.05 | 0 | SCCNV | 0.128 | 0.607 | 0.211 |
| 0.05 | 0 | SCICoNE | 0.191 | 0.299 | 0.233 |
| 0.05 | 0.05 | MintCNA | 0.750 | 0.555 | 0.638 |
| 0.05 | 0.05 | $MintCNA^{DNA}$ | 0.756 | 0.532 | 0.624 |
| 0.05 | 0.05 | rcCAE | 0.341 | 0.735 | 0.466 |
| 0.05 | 0.05 | HMMcopy | 0.106 | 0.901 | 0.189 |
| 0.05 | 0.05 | SCOPE | 0.184 | 0.861 | 0.303 |
| 0.05 | 0.05 | SCONCE | 0.099 | 0.116 | 0.107 |
| 0.05 | 0.05 | SCCNV | 0.118 | 0.607 | 0.197 |
| 0.05 | 0.05 | SCICoNE | 0.462 | 0.315 | 0.374 |
| 0.05 | 0.1 | MintCNA | 0.741 | 0.556 | 0.636 |
| 0.05 | 0.1 | $MintCNA^{DNA}$ | 0.765 | 0.531 | 0.627 |
| 0.05 | 0.1 | rcCAE | 0.376 | 0.716 | 0.493 |
| 0.05 | 0.1 | HMMcopy | 0.131 | 0.881 | 0.227 |
| 0.05 | 0.1 | SCOPE | 0.169 | 0.864 | 0.283 |
| 0.05 | 0.1 | SCONCE | 0.092 | 0.134 | 0.109 |
| 0.05 | 0.1 | SCCNV | 0.100 | 0.615 | 0.172 |
| 0.05 | 0.1 | SCICoNE | 0.628 | 0.257 | 0.365 |
| 0.1 | 0 | MintCNA | 0.842 | 0.561 | 0.674 |
| 0.1 | 0 | $MintCNA^{DNA}$ | 0.830 | 0.528 | 0.646 |
| 0.1 | 0 | rcCAE | 0.520 | 0.662 | 0.583 |
| 0.1 | 0 | HMMcopy | 0.105 | 0.950 | 0.189 |
| 0.1 | 0 | SCOPE | 0.237 | 0.870 | 0.373 |
| 0.1 | 0 | SCONCE | 0.154 | 0.678 | 0.251 |
| 0.1 | 0 | SCCNV | 0.159 | 0.697 | 0.259 |
| 0.1 | 0 | SCICoNE | 0.395 | 0.358 | 0.376 |
| 0.1 | 0.05 | MintCNA | 0.817 | 0.623 | 0.707 |
| 0.1 | 0.05 | $MintCNA^{DNA}$ | 0.825 | 0.557 | 0.665 |
| 0.1 | 0.05 | rcCAE | 0.512 | 0.637 | 0.568 |
| 0.1 | 0.05 | HMMcopy | 0.119 | 0.944 | 0.211 |
| 0.1 | 0.05 | SCOPE | 0.226 | 0.865 | 0.358 |
| 0.1 | 0.05 | SCONCE | 0.146 | 0.792 | 0.247 |
| 0.1 | 0.05 | SCCNV | 0.148 | 0.699 | 0.244 |
| 0.1 | 0.05 | SCICoNE | 0.558 | 0.211 | 0.306 |
| 0.1 | 0.1 | MintCNA | 0.817 | 0.606 | 0.696 |
| 0.1 | 0.1 | $MintCNA^{DNA}$ | 0.817 | 0.556 | 0.662 |
| 0.1 | 0.1 | rcCAE | 0.522 | 0.652 | 0.580 |
| 0.1 | 0.1 | HMMcopy | 0.141 | 0.924 | 0.245 |
| 0.1 | 0.1 | SCOPE | 0.199 | 0.858 | 0.323 |
| 0.1 | 0.1 | SCONCE | 0.138 | 0.630 | 0.226 |
| 0.1 | 0.1 | SCCNV | 0.124 | 0.701 | 0.211 |
| 0.1 | 0.1 | SCICoNE | 0.708 | 0.282 | 0.403 |

| Coverage | Noise | Method | Precision | Recall | F1 |
|---|---|---|---|---|---|
| 0.2 | 0 | MintCNA | 0.926 | 0.603 | 0.730 |
| 0.2 | 0 | MintCNA$^{DNA}$ | 0.910 | 0.554 | 0.689 |
| 0.2 | 0 | rcCAE | 0.591 | 0.665 | 0.626 |
| 0.2 | 0 | HMMcopy | 0.120 | 0.937 | 0.213 |
| 0.2 | 0 | SCOPE | 0.291 | 0.853 | 0.434 |
| 0.2 | 0 | SCONCE | 0.159 | 0.813 | 0.266 |
| 0.2 | 0 | SCCNV | 0.177 | 0.679 | 0.280 |
| 0.2 | 0 | SCICoNE | 0.459 | 0.320 | 0.377 |
| 0.2 | 0.05 | MintCNA | 0.933 | 0.613 | 0.740 |
| 0.2 | 0.05 | MintCNA$^{DNA}$ | 0.921 | 0.553 | 0.691 |
| 0.2 | 0.05 | rcCAE | 0.596 | 0.703 | 0.645 |
| 0.2 | 0.05 | HMMcopy | 0.132 | 0.926 | 0.231 |
| 0.2 | 0.05 | SCOPE | 0.285 | 0.863 | 0.429 |
| 0.2 | 0.05 | SCONCE | 0.207 | 0.932 | 0.339 |
| 0.2 | 0.05 | SCCNV | 0.163 | 0.682 | 0.263 |
| 0.2 | 0.05 | SCICoNE | 0.197 | 0.430 | 0.270 |
| 0.2 | 0.1 | MintCNA | 0.936 | 0.618 | 0.744 |
| 0.2 | 0.1 | MintCNA$^{DNA}$ | 0.934 | 0.549 | 0.691 |
| 0.2 | 0.1 | rcCAE | 0.572 | 0.704 | 0.631 |
| 0.2 | 0.1 | HMMcopy | 0.156 | 0.909 | 0.266 |
| 0.2 | 0.1 | SCOPE | 0.254 | 0.865 | 0.393 |
| 0.2 | 0.1 | SCONCE | 0.165 | 0.900 | 0.279 |
| 0.2 | 0.1 | SCCNV | 0.135 | 0.691 | 0.226 |
| 0.2 | 0.1 | SCICoNE | 0.196 | 0.360 | 0.254 |

**Supplementary Table 3. Event-level detection accuracy for deletions and duplications combined across simulation scenarios.** Precision, recall and F1 score of each method under three coverage depths (0.05×, 0.1× and 0.2×) and three noise levels (0, 0.05 and 0.1), with 90 tumor cells per **scenario**, with deletions and duplications evaluated together. A predicted CNA was counted as a true positive if it was of the same type as a ground-truth CNA and overlapped at least 20% of the ground-truth event. $MintCNA^{DNA}$: the DNA-only mode of MintCNA.

| Coverage | Noise | Method | Precision | Recall | F1 |
|---|---|---|---|---|---|
| 0.05 | 0 | MintCNA | 0.712 | 0.577 | 0.637 |
| 0.05 | 0 | $MintCNA^{DNA}$ | 0.698 | 0.540 | 0.609 |
| 0.05 | 0 | rcCAE | 0.396 | 0.702 | 0.506 |
| 0.05 | 0 | HMMcopy | 0.084 | 0.921 | 0.154 |
| 0.05 | 0 | SCOPE | 0.170 | 0.859 | 0.284 |
| 0.05 | 0 | SCONCE | 0.144 | 0.119 | 0.130 |
| 0.05 | 0 | SCCNV | 0.130 | 0.620 | 0.215 |
| 0.05 | 0 | SCICoNE | 0.218 | 0.238 | 0.228 |
| 0.05 | 0.05 | MintCNA | 0.732 | 0.568 | 0.640 |
| 0.05 | 0.05 | $MintCNA^{DNA}$ | 0.726 | 0.526 | 0.610 |
| 0.05 | 0.05 | rcCAE | 0.347 | 0.758 | 0.476 |
| 0.05 | 0.05 | HMMcopy | 0.098 | 0.908 | 0.177 |
| 0.05 | 0.05 | SCOPE | 0.166 | 0.857 | 0.279 |
| 0.05 | 0.05 | SCONCE | 0.149 | 0.127 | 0.137 |
| 0.05 | 0.05 | SCCNV | 0.123 | 0.620 | 0.205 |
| 0.05 | 0.05 | SCICoNE | 0.407 | 0.254 | 0.313 |
| 0.05 | 0.1 | MintCNA | 0.717 | 0.580 | 0.641 |
| 0.05 | 0.1 | $MintCNA^{DNA}$ | 0.727 | 0.528 | 0.612 |
| 0.05 | 0.1 | rcCAE | 0.387 | 0.724 | 0.505 |
| 0.05 | 0.1 | HMMcopy | 0.120 | 0.889 | 0.211 |
| 0.05 | 0.1 | SCOPE | 0.158 | 0.849 | 0.266 |
| 0.05 | 0.1 | SCONCE | 0.138 | 0.143 | 0.140 |
| 0.05 | 0.1 | SCCNV | 0.109 | 0.622 | 0.185 |
| 0.05 | 0.1 | SCICoNE | 0.494 | 0.211 | 0.296 |
| 0.1 | 0 | MintCNA | 0.880 | 0.588 | 0.704 |
| 0.1 | 0 | $MintCNA^{DNA}$ | 0.866 | 0.553 | 0.675 |
| 0.1 | 0 | rcCAE | 0.577 | 0.646 | 0.609 |
| 0.1 | 0 | HMMcopy | 0.107 | 0.954 | 0.192 |
| 0.1 | 0 | SCOPE | 0.237 | 0.876 | 0.373 |
| 0.1 | 0 | SCONCE | 0.208 | 0.722 | 0.323 |
| 0.1 | 0 | SCCNV | 0.173 | 0.710 | 0.278 |
| 0.1 | 0 | SCICoNE | 0.366 | 0.244 | 0.293 |
| 0.1 | 0.05 | MintCNA | 0.852 | 0.643 | 0.733 |
| 0.1 | 0.05 | $MintCNA^{DNA}$ | 0.858 | 0.580 | 0.692 |
| 0.1 | 0.05 | rcCAE | 0.573 | 0.633 | 0.602 |
| 0.1 | 0.05 | HMMcopy | 0.119 | 0.948 | 0.211 |
| 0.1 | 0.05 | SCOPE | 0.230 | 0.868 | 0.364 |
| 0.1 | 0.05 | SCONCE | 0.199 | 0.826 | 0.321 |
| 0.1 | 0.05 | SCCNV | 0.164 | 0.711 | 0.267 |
| 0.1 | 0.05 | SCICoNE | 0.395 | 0.162 | 0.230 |
| 0.1 | 0.1 | MintCNA | 0.855 | 0.637 | 0.730 |
| 0.1 | 0.1 | $MintCNA^{DNA}$ | 0.854 | 0.588 | 0.697 |
| 0.1 | 0.1 | rcCAE | 0.580 | 0.638 | 0.608 |
| 0.1 | 0.1 | HMMcopy | 0.141 | 0.930 | 0.246 |
| 0.1 | 0.1 | SCOPE | 0.211 | 0.869 | 0.339 |
| 0.1 | 0.1 | SCONCE | 0.192 | 0.656 | 0.297 |
| 0.1 | 0.1 | SCCNV | 0.144 | 0.708 | 0.239 |

| Coverage | Noise | Method | Precision | Recall | F1 |
|---|---|---|---|---|---|
| 0.1 | 0.1 | SCICoNE | 0.484 | 0.211 | 0.294 |
| 0.2 | 0 | MintCNA | 0.919 | 0.628 | 0.746 |
| 0.2 | 0 | MintCNA$^{DNA}$ | 0.906 | 0.568 | 0.698 |
| 0.2 | 0 | rcCAE | 0.608 | 0.627 | 0.617 |
| 0.2 | 0 | HMMcopy | 0.109 | 0.941 | 0.196 |
| 0.2 | 0 | SCOPE | 0.232 | 0.869 | 0.367 |
| 0.2 | 0 | SCONCE | 0.185 | 0.783 | 0.299 |
| 0.2 | 0 | SCCNV | 0.175 | 0.698 | 0.280 |
| 0.2 | 0 | SCICoNE | 0.388 | 0.212 | 0.274 |
| 0.2 | 0.05 | MintCNA | 0.919 | 0.636 | 0.751 |
| 0.2 | 0.05 | MintCNA$^{DNA}$ | 0.911 | 0.562 | 0.695 |
| 0.2 | 0.05 | rcCAE | 0.591 | 0.687 | 0.635 |
| 0.2 | 0.05 | HMMcopy | 0.120 | 0.934 | 0.212 |
| 0.2 | 0.05 | SCOPE | 0.229 | 0.874 | 0.363 |
| 0.2 | 0.05 | SCONCE | 0.228 | 0.939 | 0.367 |
| 0.2 | 0.05 | SCCNV | 0.166 | 0.701 | 0.268 |
| 0.2 | 0.05 | SCICoNE | 0.206 | 0.294 | 0.242 |
| 0.2 | 0.1 | MintCNA | 0.922 | 0.646 | 0.760 |
| 0.2 | 0.1 | MintCNA$^{DNA}$ | 0.919 | 0.566 | 0.701 |
| 0.2 | 0.1 | rcCAE | 0.589 | 0.681 | 0.632 |
| 0.2 | 0.1 | HMMcopy | 0.141 | 0.921 | 0.245 |
| 0.2 | 0.1 | SCOPE | 0.210 | 0.873 | 0.339 |
| 0.2 | 0.1 | SCONCE | 0.207 | 0.908 | 0.337 |
| 0.2 | 0.1 | SCCNV | 0.145 | 0.704 | 0.241 |
| 0.2 | 0.1 | SCICoNE | 0.214 | 0.253 | 0.232 |
